# Evaluation of Bayesian Linear Regression Models as a Fine Mapping tool

**DOI:** 10.1101/2023.09.01.555889

**Authors:** Merina Shrestha, Zhonghao Bai, Tahereh Gholipourshahraki, Astrid J. Hjelholt, Sile Hu, Mads Kjølby, Palle D. Rohde, Peter Sørensen

## Abstract

Our aim was to evaluate Bayesian Linear Regression (BLR) models with BayesC and BayesR priors as a fine mapping tool and compare them to the state-of-the-art external models: FINEMAP, SuSIE-RSS, SuSIE-Inf and FINEMAP-Inf. Based on extensive simulations, we evaluated the different models based on F_1_ classification score. The different models were applied on quantitative and binary UK Biobank (UKB) phenotypes and evaluated based upon predictive accuracy and features of credible sets (CSs). We used over 533K genotyped and 6.6 million imputed single nucleotide polymorphisms (SNPs) for simulations and UKB phenotypes respectively, from over 335K UKB White British Unrelated samples. We simulated phenotypes from low (GA1) to moderate (GA2) polygenicity, heritability (***h***^**2**^) of 10% and 30%, causal SNPs (π) of 0.1% and 1% sampled genome-wide, and disease prevalence (PV) of 5% and 15%. Single marker summary statistics and in-sample linkage disequilibrium were used to fit models in regions defined by lead SNPs. BayesR improved the F_1_ score, averaged across all simulations, between 27.26% and 13.32% relative to the external models. Predictive accuracy quantified as variance explained (R^2^), averaged across all the UKB quantitative phenotypes, with BayesR was decreased by 5.32% (SuSIE-Inf) and 3.71% (FINEMAP-Inf), and was increased by 7.93% (SuSIE-RSS) and 8.3% (BayesC). Area under the receiver operating characteristic curve averaged across all the UKB binary phenotypes, with BayesR was increased between 0.40% and 0.05% relative to the external models. SuSIE-RSS and BayesR, demonstrated the highest number of CSs, with BayesC and BayesR exhibiting the smallest average median size CSs in the UKB phenotypes. The BLR models performed similar to the external models. Specifically, BayesR’s performance closely aligned with SuSIE-Inf and FINEMAP-Inf models. Collectively, our findings from both simulations and application of the models in the UKB phenotypes support that the BLR models are efficient fine mapping tools.

## Introduction

To better understand the genetic make-up of complex quantitative phenotypes and multifactorial diseases, it is essential to be able to identify the set of genetic variants that are most likely causal or linked with the causal genetic variants. Genome-wide association studies (GWAS) often identify too many genetic variants because long-range linkage disequilibrium (LD) complicates statistical inference. Hence, additional work such as statistical fine mapping is often required to refine signal from GWAS to determine the genetic variant (or variants) most likely responsible for complex phenotypes, given verified association of region (or regions) with a phenotype. This task of determining the genetics variants and quantifying the evidence of association is crucial as it is often followed by large-scale replication studies, or laboratory functional studies to gain further biological insights for potential clinical application with drug discovery, drug-repositioning in humans. Fine mapping methods usually assume presence of potential causal genetic variants in the data (1). With the concept of existence of multiple causal genetic variants in a locus various Bayesian fine mapping methods were developed (2). Bayesian fine mapping methods can quantify uncertainty of a potential causal genetic variant through posterior inclusion probability (PIP) in a model. The PIP of a SNP refers to the mean of posterior probability that the SNP is included in a model with non-zero effect, which provides evidence for that the SNP potentially is causative (3). These methods are also able to leverage knowledge about genetic architecture of complex phenotypes through prior distribution of the effects and number of the genetic variants, which helps to improve statistical power for identifying effective genetic variants (4).

In the quest for accurate identification of effects of potential causal genetic variants in complex phenotypes, various Bayesian fine mapping methods have been developed with different modeling assumptions. FINEMAP (5) for GWAS summary statistics, uses Shotgun Stochastic Search (SSS) algorithm to search for different possible causal configurations followed by focusing on the configurations with non-negligible probability. The algorithm conducts a predefined number of iterations within the space of causal configurations. In each iteration, the neighborhood of the current causal configuration is defined by configurations that result from deleting, changing, or adding a causal SNP from the current configuration. In the next iteration, a new causal configuration is sampled from the neighborhood configurations based on its posterior normalized within the neighborhood. The sum of single effects (SuSIE) (6), a new formulation of Bayesian Variable Selection Regression (BVSR) uses a procedure called as iterative Bayesian stepwise selection approach (IBSS) to fit a model assuming few sparse causal genetic variants in a locus. It estimates the vector of regression coefficients for sparse genetic variants by summing up multiple single-effect vectors that each have one non-zero entry for a potential causal variant. Recently, (7) extended SuSIE by implementing the use of GWAS summary data and introduced it as SuSIE-RSS. SuSIE-Inf and FINEMAP-Inf (4), an infinitesimal model similar to linear mixed models, are extensions of SuSIE and FINEMAP respectively where infinitesimal effects for genetic variants in LD (estimated separately for locus) with those of the sparse components are modeled. As in FINEMAP, FINEMAP-Inf used SSS algorithm, where the posterior inference of the sparse genetic variants is marginalized over the infinitesimal effects and the residuals. As in SuSIE, SuSIE-Inf estimated the sparse causal effects by summing up multiple single-effect vectors, where for the posterior inference of sparse genetic effects the joint distribution of single-effect vectors is marginalized over the infinitesimal effects and residuals.

Following a similar pursuit of accurate fine mapping, we were interested in investigating the performance of Bayesian Linear Regression (BLR) models for fine mapping. The BLR models have been applied for mapping genetic variants, prediction (polygenic scores), estimation of genetic parameters and genetic architectures (8). The genetic architecture of a trait encompasses number, frequency and effect size of causal variants (9). The BLR models allow joint estimation of marker effects while accounting for LD among SNPs capturing the amount of variation at a genetic locus whose extent is based on the extent of the colocalization of multiple causal genetic variants. Assumptions of number and effect sizes of genetic variants are based on different prior distributions. Here we focused on the performance of the BLR models with the priors BayesC (10), and BayesR (11). During joint marker estimation, depending on priors, BLR models either shrink non-causal genetic variants’ effects or induce both variable selection method and shrinkage helping to obtain accurate estimates of effects of genetic variants. To our knowledge, the BLR models have been investigated numerous times in predictions (12-14) but only few studies have investigated precision and power of these models in fine mapping approach (4, 7).

In fine mapping, no single genetic variant is identified as causal due to the complex LD patterns between the genetic variants hence “credible sets” of potentially causal genetic variants are prioritized (1). Credible sets help in variable selection by narrowing down a larger number of variants to a small set of most likely causal variants with certain probability, refining the fine mapping approach. (15) developed a standard Bayesian approach for fine mapping, assumed a single causal locus per genetic region, to prioritize for example 99% credible set (a set whose cumulative sum of PIPs exceeds 0.99 threshold) of potentially causal genetic variants providing a credible set for per region (2). With the main goal of improving fine mapping resolution, the credible sets are meant to contain as few genetic variants as possible while still capturing an effective genetic variant.

In our study, we aimed to assess the efficiency of the BLR models (BayesC and BayesR) as a fine mapping tool using GWAS summary statistics. Using simulations, we designed credible sets and investigated precision and power of inclusion of casual variants in the credible sets to calculate *F*_1_ classification score (*F*_1_ score). We also evaluated these models based on the features of credible sets, and the prediction accuracy using fine mapped regions for five binary and five quantitative phenotypes from the UK Biobank (UKB) (16). The results from BayesC and BayesR priors were compared to the state-of-the-art methods such as FINEMAP (5), SuSIE-RSS (7), SuSIE-Inf and FINEMAP-Inf (4). We also aimed to investigate validation of the BLR model through a detailed examination of the outcomes derived from its application to Type two diabetes (T2D) within the UKB phenotypes.

## Material and Methods

In our study, efficiency of different models was investigated on simulations and the UKB phenotypes. We explored efficiency of the models on complex nature of phenotypes by simulating phenotypes with low to moderate polygenic background and creating different genetic architectures utilizing different values for heritability 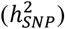, proportion of causal markers (*π*) and their effect sizes (9). Efficiency of the models was also investigated using five quantitative and five binary UKB phenotypes available from the UKB. We have discussed the theory behind single marker-linear regression analysis and its extension to summary data followed by the prior assumptions of BayesC and BayesR, used in our study. Marginal marker effects obtained from the single SNP association analysis were adjusted at multiple designed fine-mapping regions using the BLR models and external fine-mapping models. We present the design of credible sets (CSs) and definition of precision and power in terms of CSs to estimate *F*_1_ classification score (*F*_1_ score) on simulations. For the UKB phenotypes, we compared the predictive abilities (coefficient of determination: *R*^2^ for quantitative phenotypes and Area under the receiver operating characteristic curve (AUC) for binary phenotypes) and the features of CSs. Lastly, we explored the biological mechanisms underlying T2D, drawing insights from the outcomes derived by implementing the BLR model.

## Data

UKB genotyped and imputed data were used for simulations and analysis of the UKB phenotypes respectively. In our study, we had information about 488,377 participants. To obtain a genetic homogeneous study population we restricted our analyses to unrelated British Caucasians and excluded individuals with more than 5,000 missing markers or individuals with autosomal aneuploidy. Remaining (*n*=335,532) White British unrelated individuals (WBU) were used for analyses. Then, we excluded markers with minor allele frequency < 0.01, call rate < 0.95 and the markers deviating from Hardy-Weinberg equilibrium (*P*-value < 1 × 10^−12^). We excluded markers located within the major histocompatibility complex (MHC), having ambiguous allele (i.e., GC or AT), were multi-allelic or an indel (17). This resulted in a total of 533,679 single nucleotide polymorphism (SNP) markers in the simulated data. For the UKB imputed data, firstly the markers with the probability of 70% (–hard-call threshold 0.7) were converted to genotypes followed by retaining markers with imputation INFO score >= 0.8 using PLINK 2.0 (18). The same quality control criteria were applied to the imputed markers as for the genotyped data, except that we included MHC in the UKB phenotypes as this region contains many known disease-associated markers. After quality control we retained 6,627,732 SNPs and 335,532 WBU for downstream analysis in the UKB imputed data.

### Genetic architectures for simulations

To simulate genetic architectures from low to high polygenicity, we simulated quantitative phenotypes with heritability 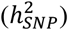 of 30% and 10%, with two different proportions of causal SNPs (*π*), 0.1% and 1%, chosen randomly from the genome.

We generated two different types of genetic architectures under a multiple regression model. In the first genetic architecture (*GA*_1_), causal SNPs (*m*_*C*_) effects were sampled from the same normal distribution:

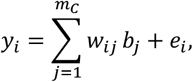

where y_*i*_ is the phenotype for *i*’th individual, *b*_*j*_ is the estimate of the *j*’th SNP effect (normally distributed with mean of 0 and variance given by 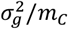). We assumed variance of a phenotype to be 1 such that 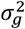 is equal to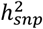. *w*_*ij*_ represents the *j*’th centered and scaled genotype of the *i*’th individual:

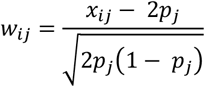

where, *x*_*ij*_ is the effect allele count for *i*’th individual at the *j*’th SNP, *p*_*j*_ is the allele frequency of the *j*’th SNP. *e*_*i*_ is the residual that has a normal distribution with mean=0 and 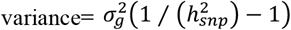. Residual variance was scaled in a way so that 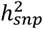 remained 30% (or 10%).

In the second genetic architecture scenario (*GA*_2_), the effects of causal SNPs are sampled from a mixture of normal distributions.

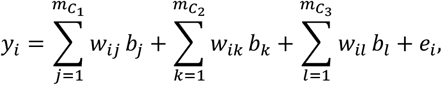

where, *b*_*j*_, *b*_*k*_, and *b*_*l*_ are the effect of causal SNPs sampled from normal distribution with mean=0 and 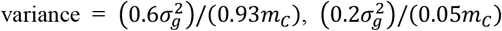 and 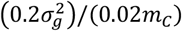 respectively. In this genetic model, the three normal distributions were designed such that 93% of the causal SNPs would have small effect sizes and the remaining 5% and 2% of the causal SNPs would have moderate and large effect sizes respectively. This genetic architecture was designed in a similar way as designed in the study by (12).

All the other parameters in *GA*_2_ are created in a similar way as for the *GA*_1_.

We created ten replicates for each simulation scenario. The total sample of 335,532 were divided into ten replicates. Each replicate contained 80% of the randomly sampled data from the total samples.

For the quantitative phenotypes, a total of eight different simulation scenarios were applied: two values of 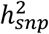, two different proportions of causal SNPs *π* and two different genetic architecture scenarios.

To simulate binary phenotypes, in addition to the parameters: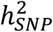, *π* and genetic architectures, we introduced another parameter “sample disease prevalence” (*PV*). Two different *PV* of 5% and 15% were used in our study. We simulated binary phenotypes from quantitative phenotypes. To simulate a binary phenotype, for example with *PV* 5%, we chose top 5% of individuals with highest simulated quantitative values as cases and the remaining as controls for the total sample in a replicate. Each scenario of a quantitative phenotype gave rise to two different scenarios for binary phenotype. In total we designed 16 different simulation scenarios for the binary phenotypes: two values of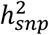, two different proportions of causal SNPs *π*, two different genetic architecture scenarios, and two prevalence *PV*. Different scenarios for the quantitative and the binary phenotypes are presented in detail in S1 Table. The flowchart of design of the simulations is presented in Fig 1.

**Fig 1.**
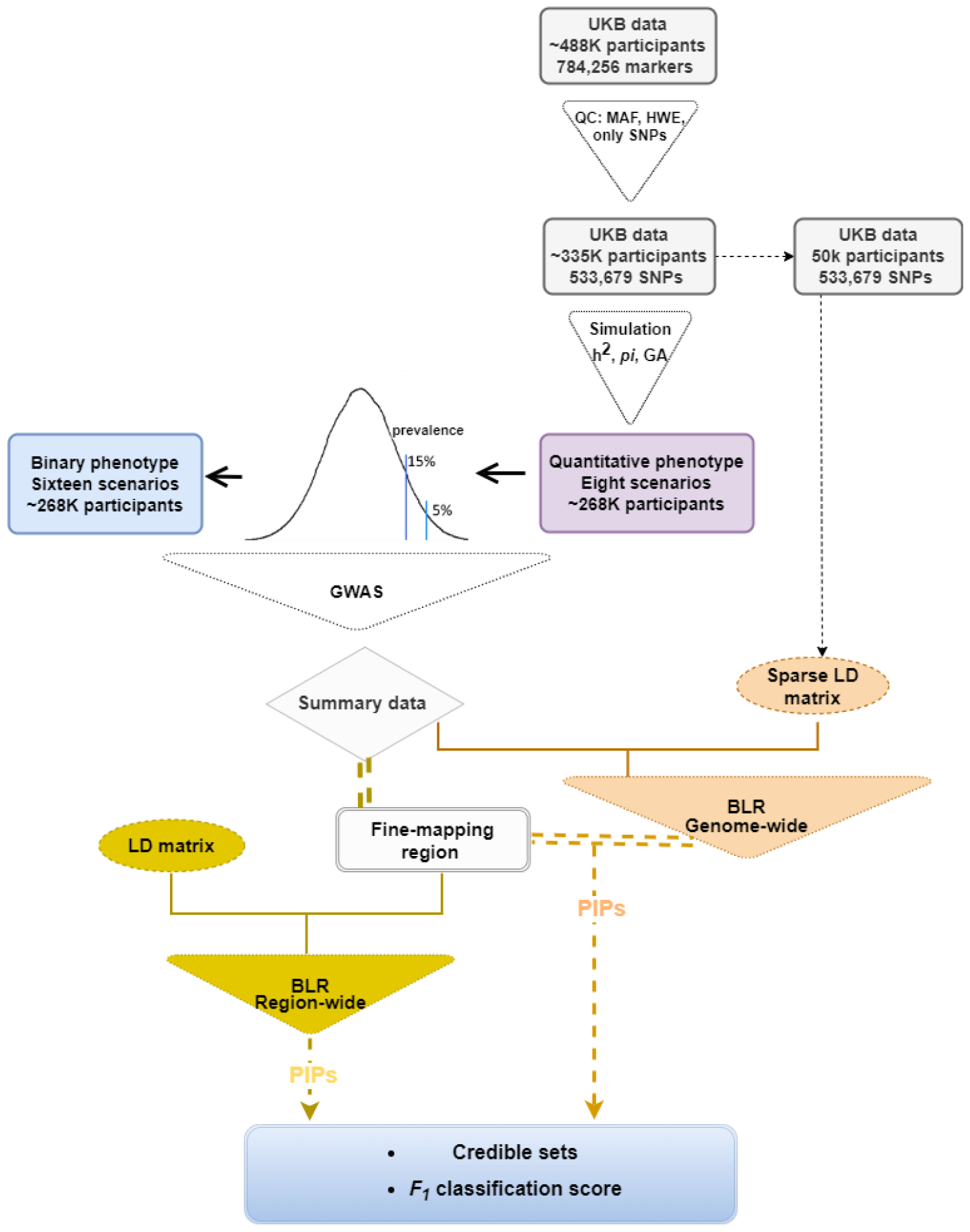
Flowchart illustrating the design of the simulation scenarios for both quantitative and the phenotypes, followed by fine mapping using Bayesian Linear Regression models. The BLR models were implemented in different ways, and the resulting posterior inclusion probability (PIPs) for SNPs were used to estimate the F_1_ classification score based on the credible sets.

### Definition of phenotypes from the UKB data

From the UKB we selected five quantitative phenotypes: Body mass index (BMI), Hip circumference (HC), Standing height (Height), Waist circumference (WC) and Waist-to-hip ratio (WHR), and five binary phenotypes: Coronary artery disease (CAD), Hypertension (HTN), Psoriasis (PSO), Rheumatoid arthritis (RA) and Type 2 Diabetes mellitus (T2D). The quantitative phenotypes were identified using specific field codes in the UKB data (see UKB showcase, Table 1a). To obtain WHR, we estimated ratio of the waist circumference to the hip circumference. In the UKB, a phenotype can have multiple instances. We used the first instance because of the least number of non-missing samples in that instance. For the definition of the binary phenotypes, to define individuals as cases for a phenotype of interest we used codes from the data field “Diagnosis-main ICD10” along with codes from the self-reported information (Table 1b). All the individuals missing the appropriate codes for the phenotype of interest were reported as controls. Additional information on age at recruitment (p21022), sex (p31), and the UKB assessment center (p54) were included as covariates in the genetic analyses. Detailed information regarding the number of samples, prevalence for the phenotypes is given in Table 1a and Table 1b.

**Table 1a.** Details of the data fields along with the total number of non-missing samples, age (mean and standard deviation), number of females (n_female) and average value for the UKB quantitative phenotypes.

| UKB Quantitative phenotypes | UKB data field | Total | Age (Mean [sd]) | Sex (n_female) | Phenotype (Mean [sd]) |
| --- | --- | --- | --- | --- | --- |
| Body Mass Index (BMI) | p21001 | 334,464 | 56.87 [7.98] | 179,309 | 27.4 [4.76] |
| Hip Circumference (HC) | p49 | 334,949 | 56.87 [7.98] | 179,517 | 103.44 [9.15] |
| Standing Height (Height) | p50 | 334,828 | 56.87 [7.98] | 179,492 | 168.86 [9.25] |
| Waist Circumference (WC) | p48 | 334,983 | 56.87 [7.98] | 179,532 | 90.37 [13.49] |
| Waist-Hip Ratio (WHR) | NA* | 334,917 | 56.87 [7.98] | 179,503 | 0.87 [0.09] |

\*NA because the phenotype was calculated in our study.

**Table 1b.** Details of the ICD10 and self-reported code used for diagnosis of cases for the UKB binary phenotypes, total number of cases, controls along with the distribution of age (mean and standard deviation) and number of females (n_female) within cases and controls.

| UKB Binary phenotypes | Definition of cases |  | Cases | Controls | Age (Mean [sd]) |  | Sex (n_female) |  |
| --- | --- | --- | --- | --- | --- | --- | --- | --- |
|  | ICD10 code [p41270] | Self-reported code |  |  | Cases | Controls | Cases | Controls |
| Coronary Artery Disease (CAD) | I21; I22; I23; I24; I25 | 1075 | 34,726 | 300,806 | 61.24 [6.38] | 56.37 [7.99] | 10,845 | 168,989 |
| Hypertension (HTN) | I10 | 1065 | 129,580 | 205,952 | 59.75 [6.98] | 55.07 [8.04] | 60,859 | 118,975 |
| Psoriasis (PSO) | L40 | 1453 | 6628 | 328,904 | 57.15 [7.88] | 56.87 [7.98] | 3090 | 176,744 |
| Rheumatoid Arthritis (RA) | M06 | 1464 | 7955 | 327,577 | 59.6 [7.04] | 56.81 [7.99] | 5251 | 174,583 |
| Type 2 Diabetes (T2D) | E11 | 1220;1223 | 25,828 | 309,704 | 60.11 [6.9] | 56.6 [8.01] | 10,072 | 169,762 |

### Statistical model

In the multiple regression model the phenotype is related to the set of genetic markers:

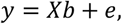

where *y* is the phenotype, *X* a matrix of genotyped markers, where values are standardized to give the ijth element as: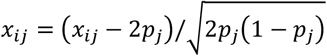, with *x*_*ij*_ the number of copies of the effect allele (e.g. 0, 1 or 2) for the ith individual at the jth marker and *p*_*j*_ the allele frequency of the effect allele. *b* are the genetic effects for each marker, and *e* the residual error. The dimensions of *y, X, b* and *e* are dependent upon the number of phenotypes, *k*, the number of markers, *m*, and the number of individuals, *n*. The residuals, *e*, are a priori assumed to be independently and identically distributed multivariate normal with null mean and covariance matrix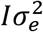.

### Extensions to summary data

The key parameter of interest in the multiple regression model is the marker effects. These can be obtained by solving an equation system like:

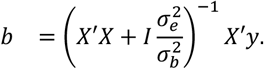

To solve this equation system individual level data (genotypes [*X*] and phenotypes [*y*]) are required. If these are not available, it is possible to reconstruct *X’y* and *X’X* from a LD correlation matrix *B* (from a population matched LD reference panel) and data (Llyod-Jones et al. 2019):

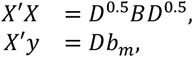

where 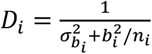 if the markers have been centered to mean 0 or *D*_*i*_ = *n*_*i*_ if the markers have been centered to mean 0 and scaled to unit variance, *b*_*i*_ is the marker effect for the i’th marker, 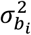 is the variance of the marginal effects from GWAS. *b*_*m*_ = *D*^−1^*X’*y is the vector of marginal marker effects obtained from a standard GWAS. The LD correlation matrix, *B*, was computed using squared Pearson’s correlation.

### Estimation of parameters using BLR models

The BLR models use an iterative algorithm, Markov Chain Monte Carlo (MCMC) gibbs sampling techniques, to estimate joint marker effects which depends on additional model parameters such as a probability of being causal (*π*), an overall marker variance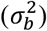, and residual variance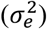. The posterior density of the model parameters 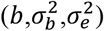 depend on the likelihood of the data given the parameters and a prior probability for the model parameters which is discussed in detail by (19).

Ideally the choice of prior for the marker effect should reflect the genetic architecture of the phenotype. Most complex phenotypes and diseases are likely highly polygenic, with hundreds to thousands of causal genetic variants, most frequently of small effect sizes (20). Thus, the prior distribution should account for many small and few large effect genetic variants. Also, marker effects are a priori assumed to be uncorrelated, but markers can be in strong linkage disequilibrium and therefore a high posterior correlation may exist. To accommodate evolving ideas genetic architectures of phenotypes and diseases, many priors for marker effects have been proposed. Each prior gives rise to a method or family of methods, and two of them are described below:

### BayesC

In the BayesC approach the marker effects, *b*, are a priori assumed to be sampled from a mixture with a point mass at zero and univariate normal distribution conditional on common marker effect variance 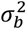. This reflects a very common thought that there were not many causal loci. This can be implemented by introducing additional variables *δ*_*i*_ which indicates if the i’th marker has an effect or not. In turn, these variables *δ* have a prior Bernoulli distribution with the probability *π* of being zero. Therefore, the hierarchy of priors is:

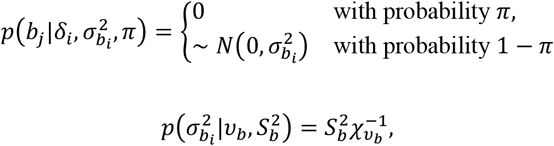

where 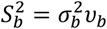 with 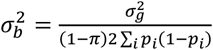 because the variance of a *t* distribution is 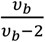.

### BayesR

In the BayesR (Erbe et al. 2012) approach the marker effects, *b*, are a priori assumed to be sampled from a mixture with a point mass at zero and univariate normal distributions conditional on common marker effect variance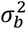, and variance scaling factors, *γ*:

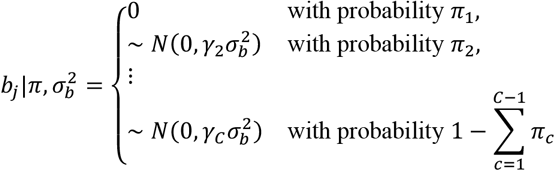

where *π* = (*π*_1_, *π*_2_, … ., *π*_*C*_) is a vector of prior probabilities and *γ* = (*γ*_1_, *γ*_2_, …. ., *γ*_*C*_) is a vector of variance scaling factors for each of C marker variance classes. The *γ* coefficients are prespecified and constrain how the common marker effect variance 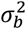 scales within each mixture distribution. Typically, *γ* = (0,0.01,0.1,1.0). and *π* = (0.95,0.02,0.02,0.01) are starting values which can be updated each iteration.

The prior distribution for the marker variance 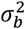 is assumed to be an inverse Chi-square prior distribution, χ^−1^(*S*_*b*_, *v*_*b*_).

The proportion of markers in each mixture class *π* follows a Direchlet (*C, c* + *α*) distribution, where c is a vector of length C that contains the counts of the number of variants in each variance class and *α* = (1,1,1,1)*’* such that the pi is updated only using information from the data.

Using the concept of data augmentation, an indicator variable *d* = (*d*_1_, *d*_2_,. ., *d*_*m*−1_, *d*_*m*_), is introduced, where *d*_*j*_ indicates whether the jth marker effect is zero or nonzero.

### Genome-wide association study (GWAS)

For simulations, we had eight and sixteen simulation scenarios (with ten replicates per scenario) for quantitative and binary phenotypes, respectively. We performed GWAS by fitting a single marker linear regression model using the R package “qgg” (19). No co-variates were used in the model because no co-variates were simulated. For analysis of the UKB phenotypes, the total population (no missing phenotype) was divided into five replicates of training (80%) and validation (20%) populations. The design for the analysis of the UKB phenotypes is presented in Fig 2. GWAS was performed in the training population of the five replicates for all the UKB phenotypes. For T2D, GWAS was also performed in the total population. We performed single marker linear regression using the R package “qgg” (19), and logistic regression analysis using PLINK 1.9 (21) for the quantitative and binary UKB phenotypes, respectively. To account for any cryptic relatedness in the data, we used top ten principal components (PCS) along with age, sex and the UKB assessment center as co-variates in the analysis of the UKB imputed data. We computed PCs for WBU from 100K randomly sampled SNPs from the genotyped data after removing SNPs in the autosomal long-range LD regions (22) with pairwise correlation (*r*^2^)>0.1 in 500Kb region, using PLINK 2.0 (18).

**Fig 2.**
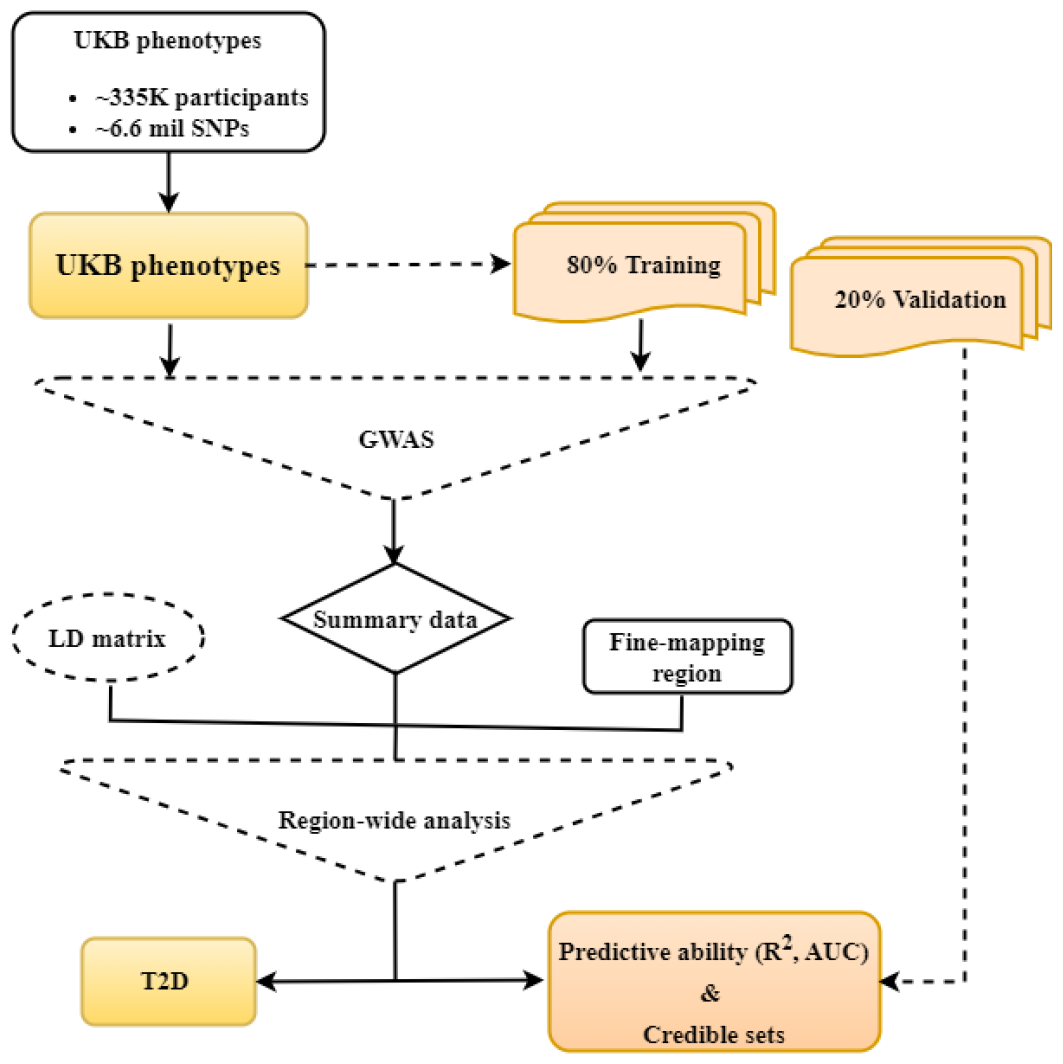
Flowchart illustrating the design of populations for the analysis of the UK Biobank phenotypes to determine the predictive abilities and features of credible sets across different models.

### Designing genomic region for fine mapping

For the simulated phenotypes, we designed fine-mapping regions based on the number of SNPs (at most 1000 SNPs in total). The regions were designed by defining a window of ∼500 SNPs to the left and right of the causal SNPs. The number of the fine-mapping regions depended on the type of simulation scenario. We did not consider any overlaps across the regions. For the UKB phenotypes, we designed the fine-mapping regions based on the physical position around each lead SNP. Significant SNPs (p-value < 5 X 10-8) from GWAS were used as the lead SNPs to design the fine-mapping regions. We defined a genomic region of one mega base pair (1MB) (∼1000kb on both sides) of the lead SNP. If the regions overlapped by more than 500kb then the regions were merged. This arbitrary number was chosen to limit the size of the regions and assuming that the SNPs added to the region might just increase the size but do not contribute to the analysis.

### Methods for fine mapping using summary statistics

We implemented BayesC and BayesR and the following external models: FINEMAP (5), SuSIE-RSS (7), SuSIE-Inf and FINEMAP-Inf (4) for fine mapping.

#### BLR models

The BLR models BayesC and BayesR, differ based on their assumption of prior variance of the marker effects. Their assumptions have already been discussed in detail above in the section “BLR models”. For the simulations, BayesC and BayesR were implemented region-wide and genome-wide, using the R package “qgg” (19). This implementation is illustrated in Fig 1. To apply these models’ region-wide, summary data from the GWAS for the SNPs in the fine-mapping regions along with the pair-wise linkage disequilibrium (LD) information among all the SNPs were used. The region-wide analysis was performed in different ways (the following three options) depending on estimation of different model parameters as part of an iterative estimation procedure (Gibbs sampling technique) from fully conditional posterior distributions. For the first option, the parameter *π* was treated as random and estimated in each iteration along with the marker variance and the residual variance. For the second option, *π* was kept constant. For the third option, only the marker variance and residual variance were estimated. Option 1: 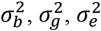 and *π* – update Option 2: 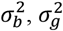 and 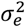– update Option 3: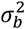, and 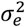– update For genome-wide application, we used summary data from the GWAS and sparse LD matrix. We randomly sampled 50,000 out of *n*=335,532 WBU to estimate sparse LD for a group of SNPs in a sliding genomic window containing 2000 SNPs, which slid 1 SNP at a time. Due to computational challenge, for genome-wide analysis, only the “Option 1” was used. The PIPs for SNPs obtained from the genome-wide analysis were used further to design credible sets for the fine-mapped regions. A total of 3000 iterations were used in the analysis with the first 500 as burn-in.

### External fine mapping tools

#### SuSIE-RSS model

The model was applied using the R package susieR (6). We provided the summary statistics (beta estimates and standard error), the LD information and number of samples for the fine mapping regions. The residual variance was estimated as suggested by the model because in-sample LD was used. We used ten causal SNPs which is the default number in the R package SusieR. We used default parameters in the functions. No priors for the SNPs were provided.

#### SuSIE-Inf and FINEMAP-Inf models

To apply these models, we downloaded python package “run_fine_mapping.py” from the link: https://github.com/FinucaneLab/fine-mapping-inf (4). We provided the summary statistics (SNP estimates and standard error) along with LD information and number of samples for the fine mapping regions. The number of causal SNPs was assumed to be ten to be consistent with the default number of causal SNPs in susieR. SuSIE-Inf and FINEMAP-Inf models were applied separately. No variance was shared and no priors for the SNPs were provided.

#### FINEMAP model

We downloaded FINEMAP software from the link: http://www.christianbenner.com/finemap_v1.4_x86_64.tgz (v1.4) (5). We provided the summary statistics (SNP estimates and standard error) along with minor allele frequency (MAF), LD information, and the number of samples for the fine mapping regions. The number of causal SNPs was assumed to be ten. No priors for the SNPs were provided.

### Quality control/convergence for models

The external fine mapping tools FINEMAP, SuSIE-RSS, SuSIE-Inf and FINEMAP-Inf, explicitly mentioned convergence of the models in the output.

For the BLR models, we estimated the convergence of the key parameters: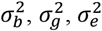, and *π*. To assess the convergence, we used the metric “zscore”. This involved calculating the difference between the average parameter values taken at the start and end of the iterations. This difference served as our metric to gauge the convergence of the desired parameter. The fine mapping regions with the absolute value of the metric “zscore”, for any of the parameters, greater than three was further investigated by thorough evaluation of the trace plots of the parameters, and scatter plots.

### Assessment of fine mapping models in simulations

Efficiency of different models were investigated based on the *F*_1_ score, a harmonic mean of precision and power estimated for the credible sets (CS).

#### Credible sets for simulations

Credible sets (CSs) help to refine association signals. The CS are defined as the minimum set of SNPs that contains all causal SNPs with probability *α*. When we assume only one causal SNP, *α* can be the sum of the PIPs for SNPs in a set. The CS in our study was designed according to (15). To design the CS of SNPs with coverage probability (cut-off or threshold) of *α*, firstly SNPs were ranked according to descending order of their PIPs. A vector of cumulative sum of PIPs was created. We added each element of the vector until it crossed a specified coverage probability of *α*. All the sets exceeding the given threshold of *α* in the fine mapping regions were refered as the CSs. A CS can contain multiple SNPs if they cross the given PIP threshold. We use a strict coverage probability of 99% for the CSs.

In simulations, with an interest to compare only the core algorithms among different models in our study, we designed the CSs for all the models irrespective of potential of the models to output the CSs. FINEMAP-Inf doesn’t give CSs, however we used the PIPs from FINEMAP-Inf to design CSs for the model. In the scenario where multiple SNPs have the same value of PIP, we investigated the list of SNPS, and if one of those SNPs is the simulated causal SNP then we included that SNP in the CS. The same procedure was applied to design CS for all the models. As we used only one causal SNP per fine-mapping regions without considering overlaps across the regions, the concept of one region harboring one causal SNP remained valid and supported our design of the CSs.

### *F*_1_ classification score for simulations

We assessed *F*_1_ score for the fine-mapping regions based on the credible sets. All the fine-mapping regions harbored a simulated causal SNP (index SNP). The *F*_1_ score takes a value between 0 and 1. The value close to 1 refers to the capability of a fine-mapping model to better identify true causal SNPs and reduce false positives.

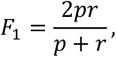

where, precision, *p* = *TP*/*TP* + *FP* and recall *r* = *TP*/*TP* + *FN*.

*F*_1_ score was calculated for each replicate of a simulation scenario. For one simulation replicate, True positive (TP) was the total number of CS (referred to as true positive CS; TP_CS) which contained the index SNP corresponding to that region. False positive (FP) was the total number of CS which crossed the threshold of alpha but did not contain the index SNP (referred to as false positives CS; FP_CS). False negative (FN) was the total number of genomic regions where the cumulative sum of PIPs did not cross the threshold of alpha, and no CS was detected. In addition to this criterion in our study for the FN, we also considered two additional criteria. We denoted “unconverged” fine-mapping regions for any methods as FN. We also considered “TP_CS” which contained more than ten SNPs as FN because large credible sets add little to no information in search of causal variants in fine-mapping procedure. We investigated the number of SNPs in true positive credible sets (TP_CS) to investigate the efficient model and tried to have the least number of SNPs in a CS as possible. The design of CSs and estimation of *F*_1_ score is represented in Fig 3.

**Fig 3.**
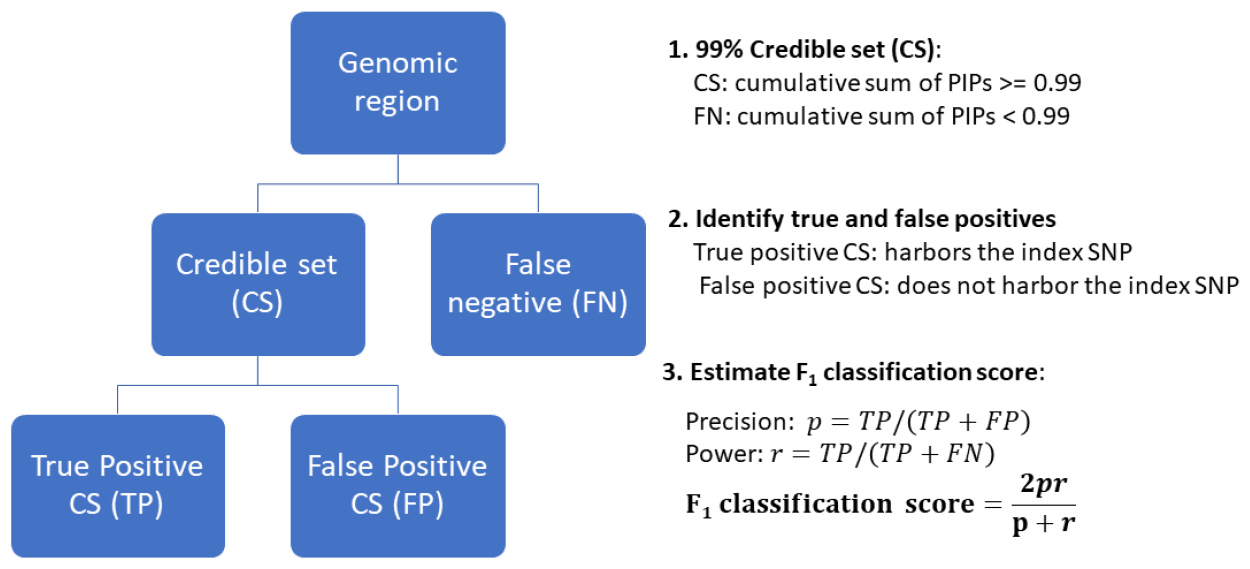
Design of credible sets with a 0.99 threshold for the cumulative sum of Posterior Inclusion Probabilities (PIPs), and estimation of the F_1_ classification score based on the credible sets.

### Investigate influence of different factors in simulations

To investigate the influence of each parameters: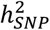, *π*, genetic architectures (*GA*) and *PV* on the performance of the models, we performed TukeyHSD test in R. To quantify the factors with the greater influence in the simulations, for each model, we also performed ANOVA on the linear model where the *F*_1_ score was regressed on 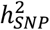, *π*, and *GA* for the quantitative phenotypes (and *PV* for the binary phenotypes).

### Similarities among model assumption in simulations

We also investigated similarities among the models based on their assumptions of genetic architectures for a complex phenotype. SuSIE, FINEMAP and BayesC assume contribution of sparse genetic variants in the genetic makeup of a complex phenotype. In addition to these sparse genetic variants, SuSIE-Inf and FINEMAP-Inf consider the influence of multiple genetic variants with small effect sizes (infinitesimal models). The BayesR model assume influence of sparse genetic variants with large effect sizes and non-sparse genetic variants with moderate to small effect sizes in the genetic makeup of a complex phenotype. We used total true positive credible sets (TP_CS) determined by each model for only the simulation scenarios for the quantitative phenotype. We investigated the number of overlaps of TP_CS of the BayesC model with SuSIE and FINEMAP, and the overlap of the BayesR with SuSIE-Inf and FIENMAP-Inf.

### Assessment of fine mapping models in the UKB phenotypes

Only the fine-mapped regions which converged across the models were used for downstream analysis to estimate predictive ability and features of the CSs.

#### Predictive Ability

For quantitative phenotypes, the predictive ability was determined by estimating the coefficient of determination, (*R*^2^). For binary phenotypes, the predictive ability was determined by estimating Area under the receiver operating characteristic curve (AUC).

Firstly, genomic score (GS) (predicted phenotype) of an individual, also known as a predictive score for a phenotype was calculated for the validation population for each replicate. GS for an individual is the sum of the product of effect alleles weighted by their estimated effect size:

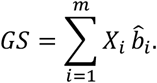

where *X*_*i*_ refers to the genotype matrix that contains an allelic count and 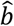 is the estimated marker effect for the *i*-th variant, *m* is the number of SNPs.

To quantify the accuracy of the GS for real quantitative phenotypes, co-variates adjusted scaled phenotypes for validation population was regressed on the predicted phenotypes. The coefficient of determination, *R*^2^, from the regression was used as a metric to assess the predictive ability of the model. To quantify the accuracy of the GS for real binary phenotypes, AUC (23)was reported:

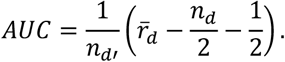

where, *n*_*d’*_ : number of controls *n*_*d*_ : number of cases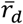: average rank of the cases.

Difference in the estimates of *R*^2^ and AUC (averaged across five replicates) among different methods was compared using TukeyHSD test.

### Credible sets for the UKB phenotypes

Unlike simulations where a fine-mapping regions were not merged irrespective of overlaps, in the UKB phenotypes, fine-mapping regions were merged if they shared a 500kb overlap of SNPs. This approach increased the likelihood of containing multiple potentially causal variants within a single fine-mapped region. To accommodate this, we designed credible sets (CSs) allowing multiple causal variants to be fine-mapped region within the same genomic region. To design a CS, in addition to the algorithm from (15) we also used information of LD. A flowchart detailing the CS design process is presented in S1 Fig. To identify significant SNP sets that are in LD, we utilized posterior PIPs and LD criteria. For each fine-mapped region, CSs were comprised of SNPs where the cumulative PIP was at least 0.80 (*PIP*_*cums*−*set*_ >= 0.80). When a CS contained multiple SNPs, the LD (r^2^) between the SNP with the highest PIP in the CS and the other SNPs was at least 0.5. Detailed steps utilized to explore the presence of multiple CSs within a fine-mapped region are mentioned in S1 Text.

We applied this methodology (S1) across all models in our study, aiming to compare the efficiency of different algorithms by using a consistent CS creation approach. This allowed us to focus solely on algorithmic efficiency by eliminating other variables. For each trait, non-converged fine-mapped regions were excluded across all the models. Afterwards, for each model, we determined the average total number of CSs, the average median CS size (SNP counts in a CS), and the average median value for the average correlations (*avg. r*^2^) among SNPs in the CS. To estimate *avg. r*^2^, we excluded the sets with only one SNP as they were not informative, and we used absolute pair-wise correlations among SNPs in the CS. In case the size of CS exceeded 100, only randomly chosen 100 SNPs were used to obtain *avg. r*^2^ for that CS. In a fine-mapped region, SNPs with *PIP*_*SNP*_ <= 0.001 was excluded before designing multiple CSs assuming that they would have little to no contribution in meeting the criterion of PIP.

### Application of BLR model in T2D

In earlier sections of our study, we examined the efficacy of the BLR models. This section delves into the application of the BLR model to a complex trait, T2D. We aimed to validate the results obtained from implementation of the BLR model.

We performed single SNP logistic regression in PLINK 1.9 (21) leveraging the entire UKB cohort for T2D (Table 1), followed by adjustments of the marginal summary statistics with the BayesR model. Fine mapping regions were created as for the UKB phenotypes discussed above in the section “Designing genomic region for fine mapping”. Multiple credible sets (CSs) per fine-mapped region were designed as discussed above in the section “Credible sets for the UKB phenotypes”.

To validate the results obtained from BayesR model for T2D, we conducted non-exhaustive comparison of our findings with the external study. Also, using the R package “gact”, we performed a gene set enrichment analysis to identify diseases enriched for T2D-associated genes and tissue-specific expression Quantitative loci (eQTLs) enrichment analysis to identify tissues enriched for T2D.

In the initial step, we mapped SNPs from multiple CSs to genes using the Ensembl Gene Annotation database available at https://ftp.ensembl.org/pub/grch37/release109/gtf/homo_sapiens/Homo_sapiens.GRCh37.87.gtf.gz. This mapping targeted SNPs within the open reading frame (ORF) of a gene, including regions 35kb upstream and 10kb downstream of the ORF, due to their potential regulatory role in controlling main ORF translation.

### Comparison with large-scale meta-GWAS study

To obtain any overlapping genes in our study with (24), one of the largest and most comprehensive meta-GWAS on T2D. The study consisted of imputed genetic variants from 898,130 European-descent individuals (9% cases). Our study limited comparison to genes given by the study in the Supplementary Table 2 which provided information of 243 loci (135 newly identified in T2D predisposition) comprising 403 unique genetic signals/associations.

**Table 2.** Average for the total number of fine mapped regions and non-converged regions for the UKB phenotypes along with the total number of credible sets, median size of the CSs, and median of the average correlations (r^2^) of the CSs for all the models.

| UKB Phenotypes | Avg. Total FMR | Avg. Non-converged FMR | BayesC |  |  | BayesR |  |  | FINEMAP |  |  | FINEMAP-Inf |  |  | SUSIE-Inf |  |  | SUSIE-RSS |  |  |
| --- | --- | --- | --- | --- | --- | --- | --- | --- | --- | --- | --- | --- | --- | --- | --- | --- | --- | --- | --- | --- |
| | | | Avg. Total CSs | Avg. Med CS size | Avg. Med $r^2$ | Avg. Total CSs | Avg. Med CS size | Avg. Med $r^2$ | Avg. Total CSs | Avg. Med CS size | Avg. Med $r^2$ | Avg. Total CSs | Avg. Med CS size | Avg. Med $r^2$ | Avg. Total CSs | Avg. Med CS size | Avg. Med $r^2$ | Avg. Total CSs | Avg. Med CS size | Avg. Med $r^2$ |
| Body Mass Index (BMI) | 219.8 | 4.8 | 310.6 | 3.4 | 0.93 | 389.8 | 2 | 0.86 | 410.6 | 5 | 0.96 | 89.8 | 8.6 | 0.97 | 204.4 | 20.6 | 0.90 | 447.2 | 15.6 | 0.95 |
| Hip Circumference (HC) | 203 | 5 | 289.4 | 1 | 0.85 | 359.4 | 1 | 0.80 | 348.4 | 2 | 0.98 | 95.6 | 3 | 0.98 | 200.6 | 4.4 | 0.97 | 410.4 | 6.8 | 0.97 |
| Standing Height (Height) | 461 | 29.6 | 1500 | 3.4 | 0.94 | 1846.8 | 2 | 0.87 | 1668 | 7.1 | 0.97 | 513.6 | 8.3 | 0.98 | 721.2 | 17.5 | 0.93 | 1696 | 14.6 | 0.96 |
| Waist Circumference (WC) | 164.4 | 2 | 229.6 | 3.6 | 0.94 | 276.8 | 2 | 0.88 | 299.4 | 6 | 0.97 | 73.2 | 7.7 | 0.98 | 157.4 | 20.8 | 0.92 | 331.6 | 15.6 | 0.96 |
| Waist-Hip Ratio (WHR) | 135.2 | 3.4 | 184.8 | 2.8 | 0.94 | 224.6 | 2 | 0.88 | 238.2 | 5.2 | 0.97 | 77 | 6.6 | 0.98 | 142.6 | 11.4 | 0.93 | 255.8 | 14 | 0.96 |
| Coronary Artery Disease (CAD) | 29.4 | 0.4 | 38.8 | 1.9 | 0.94 | 54.2 | 2.3 | 0.81 | 35.8 | 6.3 | 0.97 | 28 | 5.7 | 0.97 | 35.8 | 8.7 | 0.96 | 44 | 10.3 | 0.97 |
| Hypertension (HTN) | 137.4 | 6.2 | 211 | 2.2 | 0.90 | 277 | 2.2 | 0.80 | 239.8 | 4.2 | 0.97 | 96.8 | 5.2 | 0.98 | 159.4 | 8.8 | 0.96 | 246.8 | 9.8 | 0.96 |
| Psoriasis (PSO) | 10.8 | 3.2 | 10.2 | 3.4 | 0.92 | 9.6 | 1 | 0.89 | 9.8 | 5.3 | 0.98 | 9 | 5.7 | 0.97 | 12 | 10.5 | 0.96 | 11 | 6.4 | 0.98 |
| **Rheumatoid Arthritis (RA)** | 4 | 2.2 | 1.8 | 1.3 | NA | 1.8 | 1.2 | NA | 5.4 | 1.5 | 1.00 | 1.8 | 2.6 | 0.95 | 2.2 | 3.9 | 0.93 | 2.2 | 2.8 | 0.98 |
| **Type 2 Diabetes (T2D)** | 49.6 | 1.2 | 62.6 | 1.8 | 0.93 | 81.2 | 1.6 | 0.79 | 70.4 | 4.9 | 0.97 | 47.6 | 6.4 | 0.97 | 60 | 8.5 | 0.96 | 73.2 | 8.1 | 0.97 |

### Gene-diseases association enrichment analysis

To determine diseases significantly enriched for the gene set of our interest, we first curated a set of genes with PIP of at least 0.5 (sum of *PIP*_*SNP*_). We then downloaded the disease-gene associations data from the DISEASE database (25). This database contained disease–gene association scores (full and filtered) derived from curated knowledge databases, experiments primarily GWAS catalog, and automated text mining of biomedical literature. The analysis was conducted on the final disease-gene association data where association of a gene to a disease was combined from all the above-mentioned sources. This database includes over 10,000 diseases. However, multiple terms in the database were used to refer to the same disease. We investigated enrichment via hypergeometric test (26).

### Tissue-specific eQTLs enrichment analysis

To determine tissues enriched for eQTLs associated with T2D, firstly multi-tissue cis-eQTL annotation was obtained from GTEx (Genotype-Tissue Expression) consortium (https://storage.googleapis.com/adult-gtex/bulk-qtl/v8/single-tissue-cisqtl/GTEx_Analysis_v8_eQTL.tar) (27). We identified only eQTLs within our fine-mapped regions for each tissue. We then assessed the enrichment of tissue-specific eQTLs using a multiple linear regression model, adjusting for the influence of other tissue-specific eQTLs. The analysis was conducted using absolute beta-estimates from the BayesR model. The regression model allowed us to calculate z-scores (coefficient estimates/standard errors) and p-value for each tissue. Tissue-specific eQTLs with a p-value less than 0.05 were considered significantly enriched.

## Results

### Application in simulations

For simulations, we presented the results of the *F*_1_ score based on the credible sets to show the overall performance of the models across all simulation scenarios in Fig 4. Then to investigate the influence of each parameter considered while designing simulation scenarios, we present the results of the *F*_1_ score in each simulation scenario for the quantitative (S2 Fig) and the binary phenotypes (S3 Fig).

**Fig 4.**
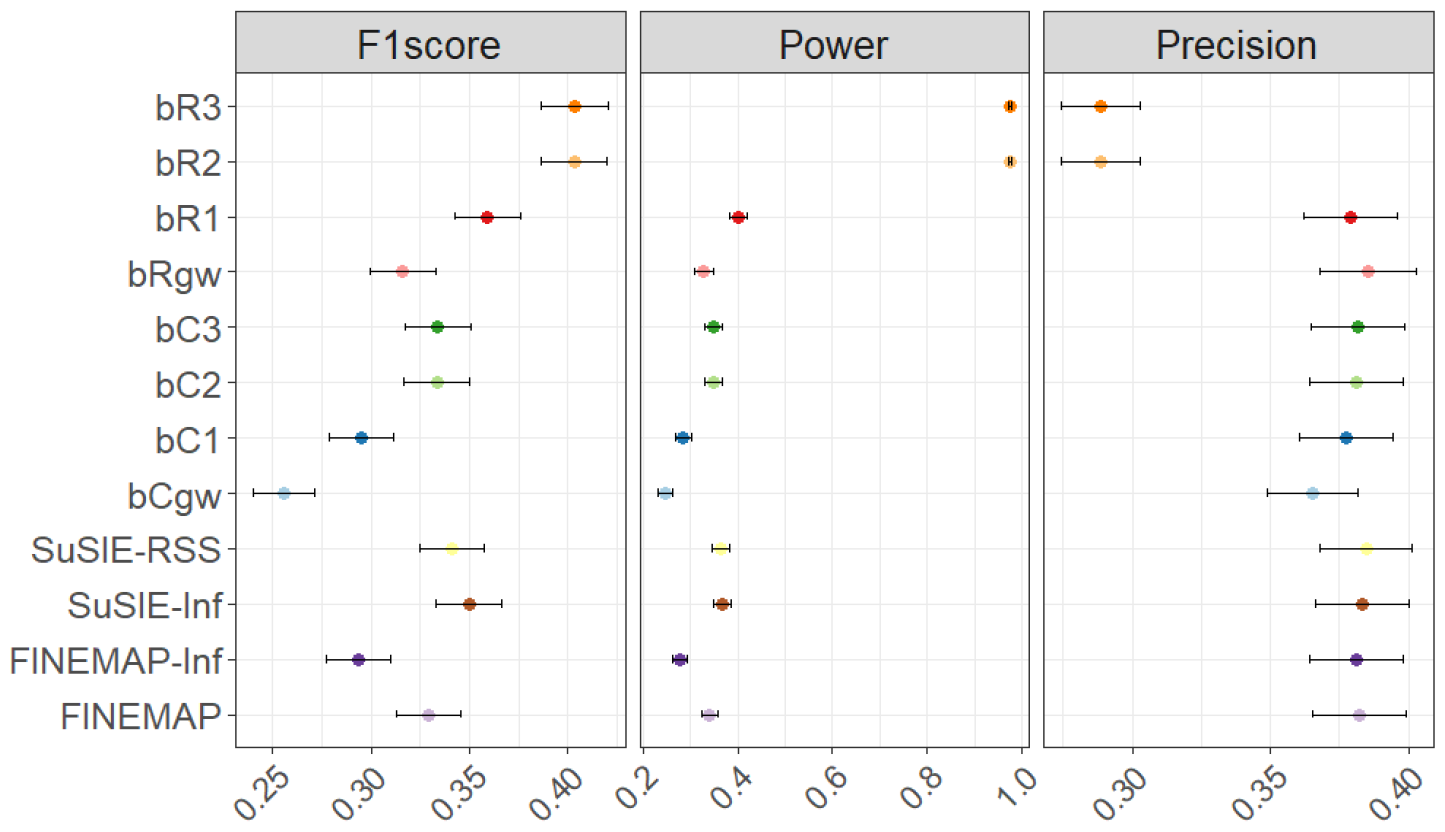
F_1_ classification score (F_1_ score), power and precision, averaged across all twenty-four simulation scenarios, for the BLR fine mapping models: BayesR region-wide models (bR3, bR2 and bR1), BayesR genome-wide model (bRgw), BayesC region-wide models (bC3, bC2 and bC1), BayesC genome-wide model (bCgw) and external models. The black solid line represents standard error for the average estimate.

### Most efficient model

The *bR*3 model (option 3) improved the *F*_1*avg,sim*_ score (average across all the simulation scenarios) by 21.64%, 11% and 0.06% relative to the BayesR genome-wide analysis (*bRgw*), and *bR*1 and *bR*2. We observed similar results for BayesC. The BayesC region-wide model (option 3, *bC*3) improved the *F*_1*avg,sim*_ score by 23.41%, 11.72% and 0.10% relative to *bCgw, bC*1, and *bC*2.

Highest *F*_1*avg,sim*_ score, averaged across all the twenty-four simulation (binary and quantitative) scenarios was observed for the BayesR region-wide model (*bR*3) [*F*_1*avg,sim*_score: 0.4] followed by SuSIE-Inf [*F*_1*avg,sim*_ score: 0.35] and SuSIE-RSS [*F*_1*avg,sim*_ score: 0.34] (Fig 1). The *bR*3 improved the *F*_1*avg,sim*_ score by 27.26%, 26.96%, 18.40%, 15.42%, and 13.32% relative to FINEMAP-Inf, *bC*3, FINEMAP, SUSIE-RSS and SUSIE-Inf. The precision (*Prec*_*avg,sim*_) and power (*Pow*_*avg,sim*_), averaged across all the simulation scenarios ranged between 0.29 to 0.39, and 0.25 to 0.98 respectively. The *bR*3 model also improved the *Pow*_*avg,sim*_ by 58% to 72% relative to other models. However, this model decreased the *Prec*_*avg,sim*_ by 26% to 33% relative to other models. Similar patterns were observed when the models were compared only within the quantitative phenotypes and within the binary phenotypes. In the following we only compared *bR*3 and *bC*3 with the external methods.

### Influence of parameters in simulations

All the models performed the best (highest *F*_1_ score) for the scenario with moderate 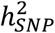 of 0.3, *π* of 0.001, and *GA*_1_ for the quantitative phenotypes (S2 Fig), and *PV* of 15% for the binary phenotypes (S3 Fig).

Pairwise comparison of the *F*_1*avg,rep*_ score (averaged across the replicates in a scenario) between all the scenarios for both the quantitative and the binary phenotypes showed significant differences between all scenarios (for all the models) as none of the intervals harbored a value of zero. S4 Fig illustrated the results for *bR3* for the quantitative simulated phenotypes. ANOVA on the results where *F*_1_ scores were regressed on 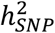, *π*, and *GA* (and *PV* for the binary phenotypes) quantified higher influence of *π* and least influence of *GA*.

### Similarities among methods assumptions in simulations

We observed that at least 50% of the true positive credible sets (TP_CS) were shared among BayesC, SuSIE-RSS and FINEMAP (S5 Fig). We observed similar results for the models BayesR, SuSIE-Inf and FINEMAP-Inf. *bCgw* identified the fewest number of total TP_CS summed across all the scenarios followed by FINEMAP-Inf. *bCgw* shared 80% of the total TP_CS with SUSIE-RSS, FINEMAP and BayesC region-wide model (*bClw* or *bC*3). The *bCgw* shared ∼91% of the total TP_CS with *bClw*. Similarly, FINEMAP-Inf shared 80% of the total TP_CS with SuSIE-Inf, *bRgw*, and *bRlw* of *bR*3. *bRgw* shared 85.1% of the total TP_CS with *bRlw* or *bR*3.

## Application to UKB phenotypes

### Predictive ability

We observed a significant decrease in the 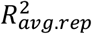 (averaged across all the binary phenotypes) of BayesC and BayesR relative to SuSIE-Inf and FINEMAP-Inf for the phenotypes BMI, WC, HC and WHR (Fig 5a). No significant difference in the 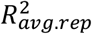 was observed between BayesR compared to SuSIE-Inf and FINEMAP-Inf for Height, whereas a significant decrease was observed for BayesC compared to these models. We observed significant improvement in the 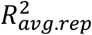 of BayesR relative to SuSIE-RSS for Height. All the methods could predict Height better compared to other quantitative phenotypes. Prediction *AUC*_*avg,bin*_ (averaged across all the binary phenotypes) with BayesR increased by 0.40%, 0.16%, 0.08%, 0.05% compared to SUSIE-RSS, BayesC, FINEMAP-Inf and SuSIE-Inf, respectively (Fig 5b). We didn’t observe any significant differences between the *AUC*_*avg,rep*_ (averaged across all the replicates) of models compared pairwise for any binary phenotypes except for HTN. For HTN, BayesR improved the *AUC*_*avg,rep*_ significantly compared to SuSIE-RSS. The highest estimate of the *AUC*_*avg,rep*_ was observed for T2D followed by HTN for all the models. The lowest estimate of the *AUC*_*avg,rep*_ was observed for RA. Prediction 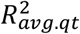 (averaged across all the quantitative phenotypes) with BayesR decreased by 5.32% and 3.71% compared to SuSIE-Inf and FINEMAP-Inf, whereas increased by 7.93% and 8.3% compared to SuSIE-RSS and BayesC. BayesR model improved the 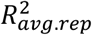 (averaged across all the replicates) significantly compared to BayesC model for all the quantitative phenotypes except for WHR.

**Fig 5.**
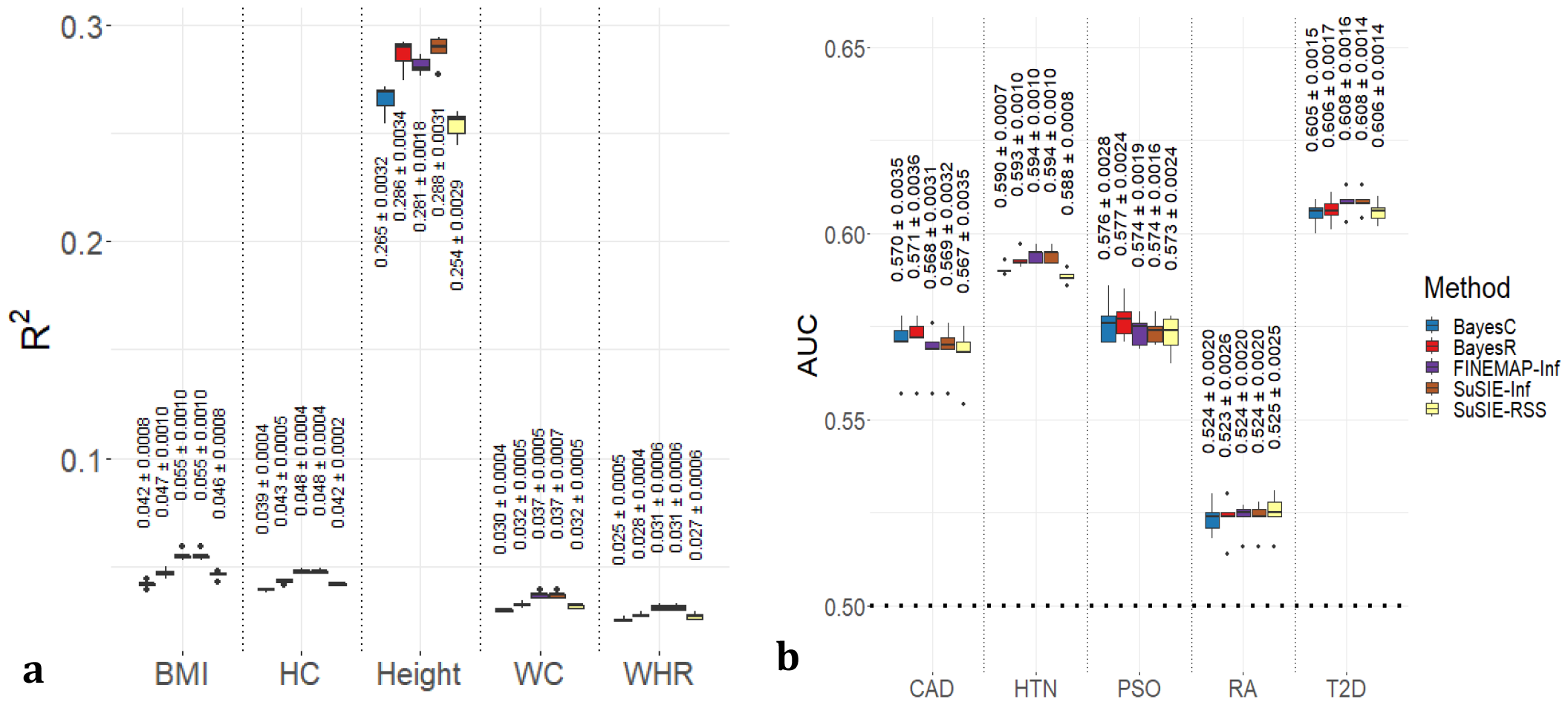
Prediction accuracies estimated from fine mapped regions. **a**. Bar plot of prediction accuracy, represented by the coefficient of determination (R^2^), averaged across five replicates for the UKB quantitative phenotypes: body mass index (BMI), hip circumference (HC), standing height (Height), waist circumference (WC), and waist-hip ratio (WHR). **b**. Bar plot of prediction accuracy, represented by the Area under the Curve (AUC), averaged across five replicates for the UKB binary phenotypes: coronary artery disease (CAD), hypertension (HTN), psoriasis (PSO), rheumatoid arthritis (RA), and type 2 diabetes (T2D). The models used in the fine mapping can be identified by the colors in the legend associated with each model. For each method within a trait, corresponding mean of R2 or AUC across five replicates and standard error is written on the top of the box plot.

### Credible sets

The average total number of fine-mapped regions across five replicates for the quantitative phenotypes ranged from 135.2 for WHR to 461 for Height and for the binary phenotypes ranged from 4 for RA to 137.4 for HTN (Table 1). The highest averaged non-converged regions were observed for RA (55%) followed by PSO (29.62%). For other phenotypes, the non-converged regions ranged from 1.22% to 6.42%.

BayesR determined the highest average number of CSs for Height, CAD, HTN and T2D, whereas SuSIE-RSS determined the highest average number of CSs for BMI, HC, WC and WHR (Table 1). For the above-mentioned phenotypes, FINEMAP-Inf determined the smallest average number of CSs. All the models obtained a similar average number of CSs for PSO (9 to 12) and RA (1.8 to 2.2).

The BLR models showed the smallest average median CS size across all the phenotypes compared to the external fine-mapping models (Table 2). BayesR showed the smallest average median size of CS for BMI, Height, WC, WHR, PSO, RA and T2D. BayesC showed the smallest average median size of CS for CAD. Both BayesC and BayesR showed the same average median size for HC and HTN. The highest average median CS size was shown by SuSIE-Inf for BMI, Height, WC, CAD, PSO, RA and T2D. For other phenotypes, SuSIE-RSS showed the highest value for the median CS size.

The average median for *avg. r*^2^ for the BLR models were smaller compared to the external models. BayesC showed the largest average median value compared to BayesR across all the phenotypes.

### Application of BLR model in T2D

We identified a total of 117 CSs for T2D across 69 fine-mapped regions with a median CS size of 2 (range:1 to 297), and the median of *avg. r*^2^ was 0.80 (range: 0.49 to 1). We identified 53 CSs of size 1 (1 SNP counts), 47 CSs of size between 2 to 50, and the remaining 17 CSs of size more than 50 SNPs.

### Comparison with large-scale meta-GWAS study

We found 53 of the 181 genes identified from our study, listed in Table S2, overlapped with genes from the study Mahajan et al. (2018) (S2 Table). Among 53 overlapped genes, 10 genes (*DTNB, RBM6, MBNL1, SLCO6A1, PDE3B, CELF1, MAP2K7, ZC3H4, EYA2*, and *ZBTB46*) were categorized as novel associations in the study by (24).

Additionally, our study identified multiple SNPs at *TCF7L2* in addition to rs7903146 (PIP: 0.9996). This includes rs34855922 (PIP: 0.3844), rs11196234 (PIP: 0.3512) and rs7912600 (PIP: 0.086) within a CS (*avg. r*^2^: 0.70), as well as rs145034729 (PIP: 0.992) linked to *TCF7L2* locus.

### Gene-Diseases association enrichment

The top 30 significant diseases (p-value < 0.05) enriched for our T2D-related gene set and their corresponding p-values are detailed in S3 Table. The list includes disease terms such as Type 2 Diabetes Mellitus, Diabetes Mellitus, ICD10:E11 code for T2D, as used in the UKB database. Additionally, we discovered associations with various forms of diabetes, such as several types of maturity-onset diabetes of the young (MODY), prediabetes syndrome, gestational diabetes, both permanent and transient neonatal diabetes, ICD10-E14 (unspecified T2D), and ICD10-O24 (diabetes in pregnancy). The list also encompassed other conditions, including Rheumatoid Arthritis (RA) with corresponding ICD10 codes: M0, M05, M06 and M069, Wolfram syndrome, hyperglycemia, hyperinsulinism, glucose intolerance, pancreatic agenesis, pancreatic cystadenoma, and insulinoma.

### Tissue-specific eQTLs enrichment

Among 49 different tissues, significant enrichment (p-value < 0.05) of T2D-related eQTLs were identified in the 13 tissues (S6 Fig): Brain cerebellar hemisphere (n=419), Cells cultured fibroblasts (n=718), Brain cerebellum (n=467), Pituitary (n=379), Esophagus muscularis (n=615), Brain nucleus accumbens basal ganglia (n=308), Lung (n=624), Skin (not sun exposed suprapubic) (n=678), Artery tibial (n=647), Adipose subcutaneous tissue (n=695), Muscle skeletal tissue (n=639), Thyroid (n=810), and Nerve Tibial (n=804).

## Discussion

Here we aimed to assess the efficiency of BayesC and BayesR as a fine mapping tool. We applied these models in simulations and the real UKB data using summary statistics. In simulations, the efficiency was investigated based on *F*_1_ score. For the UKB phenotypes the models efficiency was based on polygenic scores and credible sets. BayesC and BayesR models’ efficiency were compared to the state-of-the-art methods such as FINEMAP (5), SuSIE-RSS (7), SuSIE-Inf and FINEMAP-Inf (4). All the models used in our study serve the same purpose of identifying true effects of causal variants. However, they differ in the details in the algorithm and their implementation which applied together can have different impact on the overall performance.

### BayesC and BayesR

BayesC and BayesR applied genome-wide and region-wide have the same assumptions of prior variance of marker effects, but they differed in their implementation in our study. To our knowledge this is the first study comparing implementation of BayesC and BayesR in such manner. We implemented the models, genome-wide where the posterior distributions of the model parameters were estimated based on taking SNPs genome-wide whereas region-wide implementation were limited to the fine-mapped regions designed based on the simulated causal SNPs. In the simulations, better performance of both priors when implemented region-wide compared to genome-wide based on the *F*_1*avg,sim*_ score for both BayesC and BayesR. However, the genome-wide models showed better *Prec*_*avg,sim*_ but less *Pow*_*avg,sim*_ (fewer true positive CSs) than region-wide models. High percentage of overlaps of the CSs for the genome-wide models with the region-wide models suggests that there is a potential in the genome-wide models. It would be interesting to investigate further the common CSs determined by the genome-wide and the region-wide models.

BayesR showed significant improvement in prediction accuracy, *Prec*_*avg,rep*_ for four out of the five quantitative phenotypes relative to BayesC. Our prediction accuracies are consistent with previous studies (28, 29). (28) showed an increase of prediction ability, averaged across various economical phenotypes in cattle, using BayesR compared to BayesC. (29) showed similar results across simulation scenarios for phenotypes with high heritability. BayesR identified a large number of CSs, and small sized CSs relative to BayesC. Our results suggest that BayesR assumption about genetic architecture suits better for polygenic phenotypes predictions where many different effect sizes are observed, relative to BayesC

### Comparison of the BLR models to external models

To our knowledge this is the first study comparing BayesR model to the state-of-the-art models: FINEMAP, SuSIE-RSS, SuSIE-Inf and FINEMAP-Inf. Across different simulation scenarios, BayesR had higher *F*_1*avg,sim*_ score relative to the external models with high power but with less precision. The average prediction accuracy 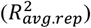 for the quantitative phenotypes was significantly lower for BayesR and BayesC compared to SuSIE-Inf and FINEMAP-Inf for BMI, HC, WC and WHR. We observed highest estimates for the predictive ability of the infinitesimal models, which might have been because we used SNP effects from both sparse and infinitesimal components for SuSIE-Inf and FINEMAP-Inf for predictions, unlike in the study (4) where only sparse components were used to compare prediction accuracies between SuSIE, FINEMAP, SuSIE-Inf and FINEMAP-Inf. Since infinitesimal components do not neglect any SNP effects, this may also explain high prediction accuracy for the models including infinitesimal effects. Our results showed that the performance of BayesR is closer to the infinitesimal models.

(30) compared the performance of BayesC to various methods including SuSIE and FINEMAP in fine mapping, where BayesC performed similar to SuSIE but better than FINEMAP in power and false discovery rate determination, for different simulation scenarios. We showed that BayesC had improved power relative to FINEMAP whereas the power was decreased relative to SuSIE-RSS. This difference in results might be due to differences in implementation of these models as this study applied the models in whole-genome scale using local regression approach where we applied the models only in specific regions defined by simulated causal SNPs, that not necessarily included whole genome. Our study compared SuSIE-RSS (which is an extension of SuSIE that uses summary statistics) and BayesC, FINEMAP using summary statistics with in-sample LD among other models, whereas this study used individual levels data for BayesC and SuSIE, and summary statistics with in-sample LD for FINEMAP.

BayesR and SuSIE-RSS identified a greater number of CSs when applied to the UKB phenotypes. However, BayesR showed the smallest average median CS size. We constructed multiple credible sets for all the models based on our algorithm where we applied the cut-off thresholds of 0.80 for a set to be a CS. We are aware that FINEMAP, SuSIE-RSS and SuSIE-Inf also determine CS where multiple CS can be determined based on multiple causal variants in the fine mapped region. Such procedure of determining CSs might alter the features of the CSs. However, the main objective of our study was to compare the efficiency of the algorithms of these models. Introducing a comparison based on the CSs they determine would introduce additional complexity and divert us far from our objective. Hence, we determined multiple CSs using the same algorithm for all the models. However, it would be interesting to compare CSs developed by the external models and evaluate their efficiency.

### Influence of parameters in simulations

We observed a significant difference in F_1_ score between the different simulated parameters 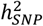 (30% and 10%), *π* (0.001 and 0.1) and *GA* (*GA*_1_ and *GA*_2_), and *PV* (5% and 15%). The pairwise comparison of *F*_1*avg,rep*_ score, within a scenario, among different simulation scenarios for each model showed significant differences among scenarios and significant contribution of each parameter. However, the large value of the F-statistic obtained from ANOVA on the results of the regression 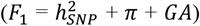 was seen for the parameter *π* suggesting greater influence of this parameter in performance of the model. In our simulation, a smaller number of causal SNPs for a given genetic variance would be sampled from a larger marker effect variance compared to a higher number of causal SNPs. This large effect SNPs must have high PIPs such that the credible sets determined by the models harbored the true causal SNPs. The *F*_1_ score was based on the detection of a true simulated causal variant in a credible set. In addition to the threshold for a cumulative sum of PIPs [0.99] that a set needs to cross to be a credible set, we also set a limit on the size of CS (not more than 10). The main motive of the CS was to refine the resolution of the fine mapping region and a CS with large number of SNPs even if it harbored a true causal variant would not be informative. We used a strict cut-off threshold of 0.99 for cumulative sum of PIPs and maximum size of 10 for CS. The results might differ with lenient thresholds for the cumulative sum, and size of CS. As per our expectation, all the models performed significantly better (high *F*_1*avg,rep*_ score) for the phenotypes with moderate 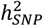 [30%], a smaller number of causal SNPs [*π*: 0.001], and the phenotypes simulated with few SNPs with large effect size [*GA*_1_] for the quantitative phenotypes, and the worst performance was observed for the phenotypes with low 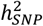 [10%] and a larger number of causal SNPs [*π*: 0.01].

### *F*1 classification score – power and precision of the BLR models

F_1_ score is a harmonic mean of precision and power/recall and is a well-known performance metric used for model comparison especially under class imbalance. It penalizes the performance even when only one of either precision or power is low. In our study, both precision and power are given equal importance for the performance of a model. We observed higher *Pow*_*avg,sim*_ for BayesR compared to other models. Highest power of BayesR referred to the scenario where majority of CSs obtained from BayesR had small size CSs. We used in-sample LD, while using external summary statistics in-sample LD is not always available as also mentioned by (30). Hence, the power may decrease while using an external reference LD panel. We observed low *Prec*_*avg,sim*_ of BayesR. This referred to as substantial amount of CSs were false positives. The range of *Prec*_*avg,sim*_ across all the models is not vast suggesting that all the models showed similar performance for precision.

### The UKB phenotypes, accuracy and fine mapping, credible sets

The predictive accuracies for the UKB phenotypes were smaller compared to other studies. 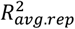 for BMI, Height, HC, WHR, and *AUC*_*avg,rep*_ for T2D for the UKB data presented by (12) using SbayesR and around 1.1 million SNPs were larger compared to the values estimated using BayesR in our study. In our study, accuracies were derived from imputed SNPs limited only to the fine mapped regions. For polygenic phenotypes for example in Height and BMI, (31) suggested enrichment of heritability from rare genetic variants (MAF < 0.01). In our study, we discarded rare SNPs with MAF < 0.01 and focused only on common SNP effects. In our study, non-converged regions for a model were excluded from analysis for prediction accuracies (also credible sets) which might also have impacted the estimated accuracies.

### Validation of BLR model

Compared to the recent meta-GWAS on T2D (24), we identified 10 genes to overlap with the 53 genes, which were categorized as novel loci in (24). This finding demonstrates the effectiveness of BayesR model combined with credible sets in identifying potential causal variants, even in studies with comparatively smaller size. This limited number (53) of overlapping genes could be attributed to our study’s smaller scale (25,828 cases and 309,704 controls compared to 74,124 cases and 824,006 controls in (24)), which could limit ability to detect especially rare variants, and the exclusion of rare variants (excluding SNPs with < 1% MAF in our study). Additionally, the discrepancies in how SNPs were mapped to a gene between our study and that of (24) might also contribute to this limited overlap.

*TCF7L2* (Transcription Factor 7-like 2) explained the highest genetic variance (0.035) in our study. This gene plays a crucial role in Wnt signaling pathway, which regulates pancreatic islet cell proliferation and survival (32). In *TCF7L2*, rs7903146 is the largest-effect common variant signal for T2D in Europeans (24). Observation of multiple signals for T2D at *TCF7L2* in addition to rs7903146 in (24) was the first evidence according to this study. In addition to the rs7903146, we also identified SNP rs34855922 associated to T2D similar with (24), which again demonstrates the effectiveness of BayesR model combined with CSs. The rs7903146 and rs34855922 are two of the eight SNPs that mark regulatory elements within *TCF7L2* locus (33). The rs7903146 coordinate regulation of *TCF7L2* expression, and overlaps histone modification marks and an annotated enhancer in the pancreas (33). Our study also identified an intronic variant (rs145034729) at the *TCF7L2* locus. The effect of this intronic SNP is uncertain. However, it may function as an enhancer element, modulating the expression of distal genes without necessarily affecting the function of *TCF7L2* itself. The discovery of multiple variants within the *TCF7L2* locus is interesting, as (33) suggests that it acts as a regulatory hub for genes implicated in the etiology of T2D. Identifying these variants in this locus offers valuable insights into the biological mechanisms underlying T2D.

The gene set enrichment analysis for diseases provided further support for the efficacy of BayesR model in T2D. This analysis revealed significant enrichment of our gene set for diseases such as T2D, hyperglycemia (diabetes-like symptoms), hyperinsulinism (one of the processes leading to hyperglycemia (34). Significant enrichment to other types of diabetes and diseases may reflect shared genetic factors (via pleiotropic genes or common pathways) influencing the etiology of diverse conditions (diseases) through different mechanisms. For instance, (35), noted an increased risk of diabetes mellitus incidence in individuals with RA, highlighting the potential role of inflammatory pathways in the T2D pathogenesis.

For tissue enrichment analysis, our findings indicate that T2D related eQTLs exhibit tissue-specific effects on gene expression. The implications of our results can be viewed from multiple perspectives. Our results may suggest a complex interplay of regulatory regions in significantly enriched tissues leading to T2D predisposition. Our results may also suggest individuals with T2D might experience adverse effects in these tissues, potentially leading to a range of complications. For instance, (36) explored the association of significantly enriched tissue specific T2D associated eQTLs with different T2D complications. Here we delve into the cerebellar hemisphere region of the brain, the most significant enriched tissue. This region, part of the cerebellum (another significant tissue in our study), has been linked to cognitive impairments when abnormal. (37) highlighted significant cognitive impairments in T2D individuals, correlating these deficits with considerable loss in gray matter volume in brain regions associated with these functions. The decline in insulin transport and resistance in the cerebral cortex, an area dense with high insulin receptor, may impair regional glucose metabolisms, leading to gray matter volume changes potentially leading to structural and functional changes in brain in T2D individuals.

No association with pancreatic tissue was found, likely due to the GTEx database’s limitations. The pancreatic tissue in GTEx represents mostly (97%) exocrine cells that mask islets signals (38). Pancreatic islets are clusters of specialized endocrine cells that are essential to maintain glucose homeostasis and play a central role in etiology of T2D.

Our study was confined to the cis-eQTLs database from GTEx consortium. (39) have shown that trans-eQTLs contribute significantly to T2D heritability, suggesting that further exploration of trans-eQTLs could enhance the understanding of gene expression and cellular functions across different tissues.

In conclusion, we observed that the performance of the BLR models was comparable to the state-of-the-art external models. The performance of BayesR prior was closely aligned with SuSIE-Inf and FINEMAP-Inf models. Results from both simulations and application of the models in the UKB phenotypes suggest that the BLR models are efficient fine mapping tools.

## Data availability statement

The genetic and phenotypic data utilized in our study were obtained from the UK Biobank Resource (ID 96479).

## Ethics statement

Human studies in the UK Biobank project have received approval from the Ethics and Governance Framework (EGF), which ensures data and sample usage adheres to scientific and ethical standards. The consent to participation will apply throughout the lifetime of the UK Biobank, unless participants withdraw, and involves the collection and storage of biological samples (blood, saliva, urine) and electronic health records (GP, hospitals, dental, prescriptions). Individual data is anonymized, with each research project receiving its own anonymized dataset. The ethics committee waived the need for written informed consent.

## Author Approvals

All authors have seen and approved the manuscript, and it hasn’t been accepted or published elsewhere.

