## Supplementary Figures S1-S6 for "Evaluation of Bayesian Linear Regression Models as a Fine Mapping tool"

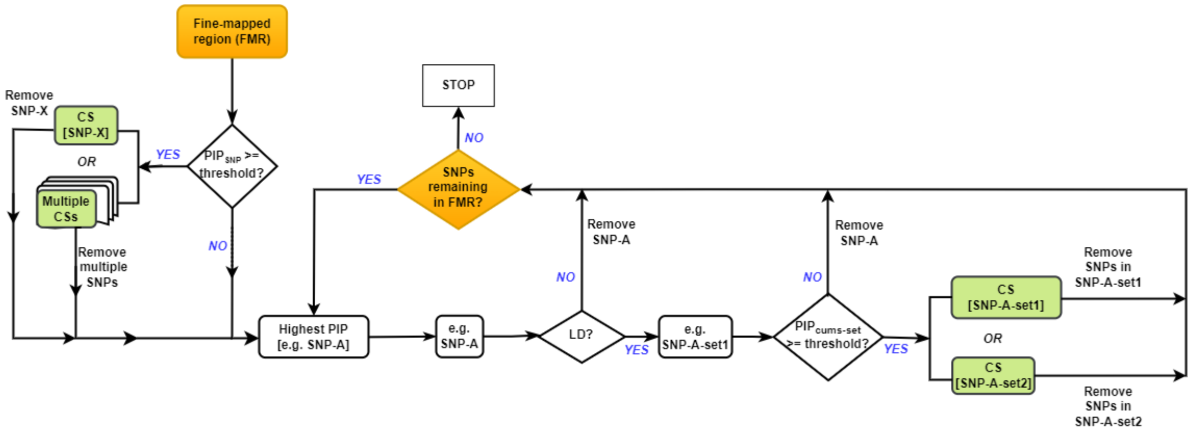


**S1 Fig. Flowchart illustrating methodology for defining multiple credible sets (CSs) in relation to the UK Biobank (UKB) phenotypes.** After establishing criteria or thresholds for the posterior inclusion probability (PIP) and linkage disequilibrium (LD) for the set of SNPs, in a fine mapping region, any SNPs (for example SNP-X) meeting the PIP criteria formed a CS. Multiple SNPs meeting this criterion formed multiple CSs. SNP/SNPs in CS/CSs were excluded. The analysis continued with the remaining SNPs or if none of the SNPs met the PIP criteria. Then a SNP with the highest PIP (SNP-A) is recognized followed by identifying SNPs (SNP-A-set1), in LD with SNP-A, that met LD criteria. In the absence of SNP-A-set1, SNP-A was removed from the fine mapping region without forming any CS. If SNP-A-set1 exists, the cumulative sum of descending order of the set was evaluated against the PIP criteria. If the set failed the PIP criteria, SNP-A was removed. If the set met the PIP criteria, the set was recognized as a CS, and the SNPs in the set were excluded from the fine mapping region. The resulting CS might be either the set SNP-A-set1 or a subset thereof, like SNP-A-set2. This process was iterated until no SNPs remained for analysis in the fine-mapped region.

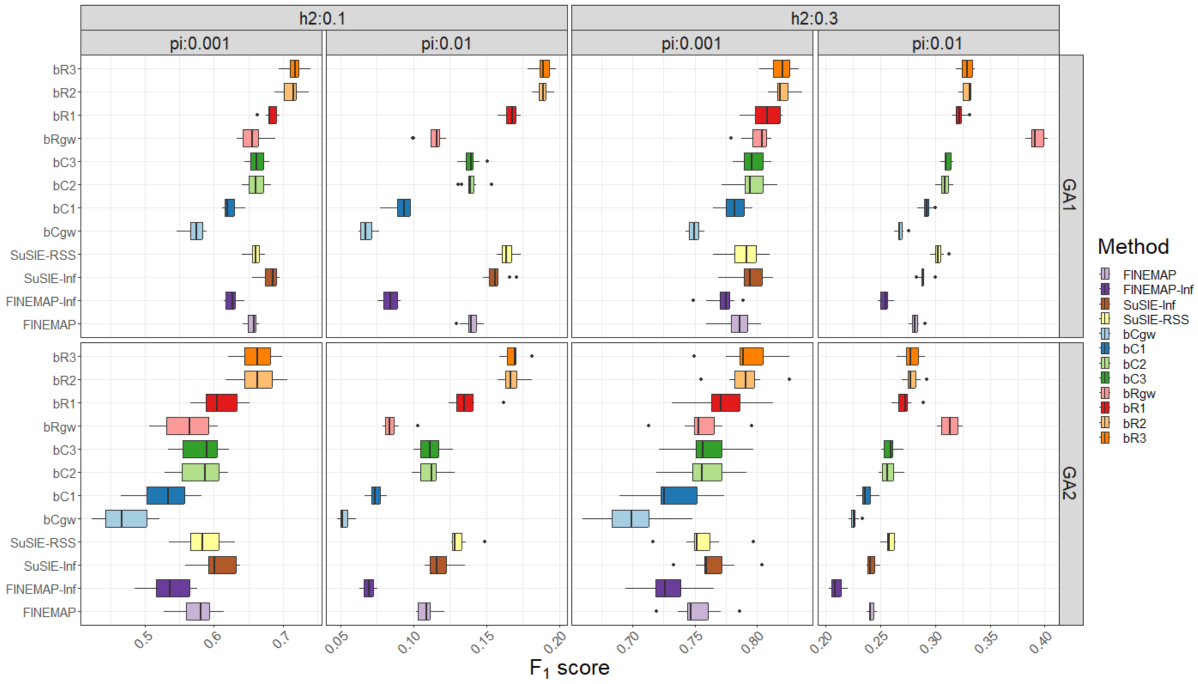


**S2 Fig. Box plot for $F_1$ scores, averaged across ten replicates, for eight simulation scenarios, which encompass heritabilities ($h^2$) of 0.3 and 0.1, proportion of causal genetic variants ($\pi$) of 0.001 and 0.01, and genetic architectures GA1 and GA2.** These scenarios pertain to simulated quantitative phenotypes. $F_1$ scores were estimated from credible sets for the BLR fine mapping models: BayesR region-wide models (bR3, bR2 and bR1), BayesR genome-wide model (bRgw), BayesC region-wide models (bC3, bC2 and bC1), BayesC genome-wide model (bCgw) and external models.

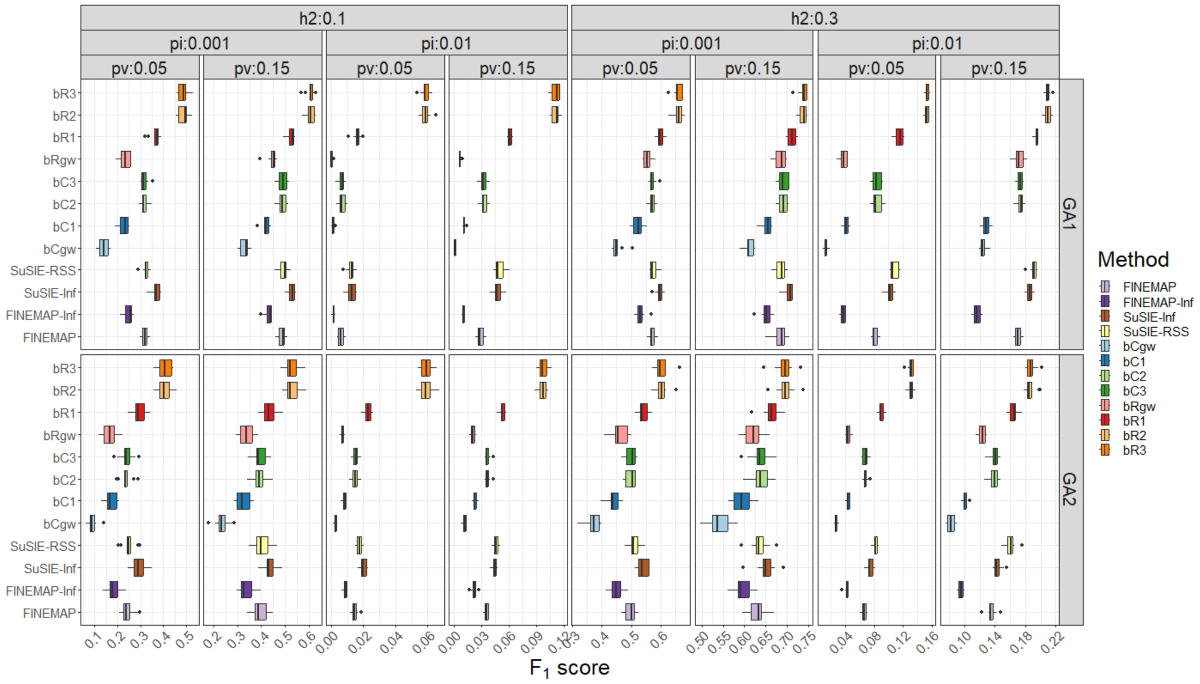


**S3 Fig. Box plot for $F_1$ scores, averaged across ten replicates, for sixteen simulation scenarios, which encompass heritabilities ($h^2$) of 0.1 and 0.3, proportion of causal genetic variants ($\pi$) of 0.001 and 0.01, genetic architectures GA1 and GA2, and prevalence of 0.5 and 0.15.** These scenarios pertain to simulated binary phenotypes. $F_1$ scores were estimated from credible sets for the BLR fine mapping models: BayesR region-wide models (bR3, bR2 and bR1), BayesR genome-wide model (bRgw), BayesC region-wide models (bC3, bC2 and bC1), BayesC genome-wide model (bCgw) and external models.

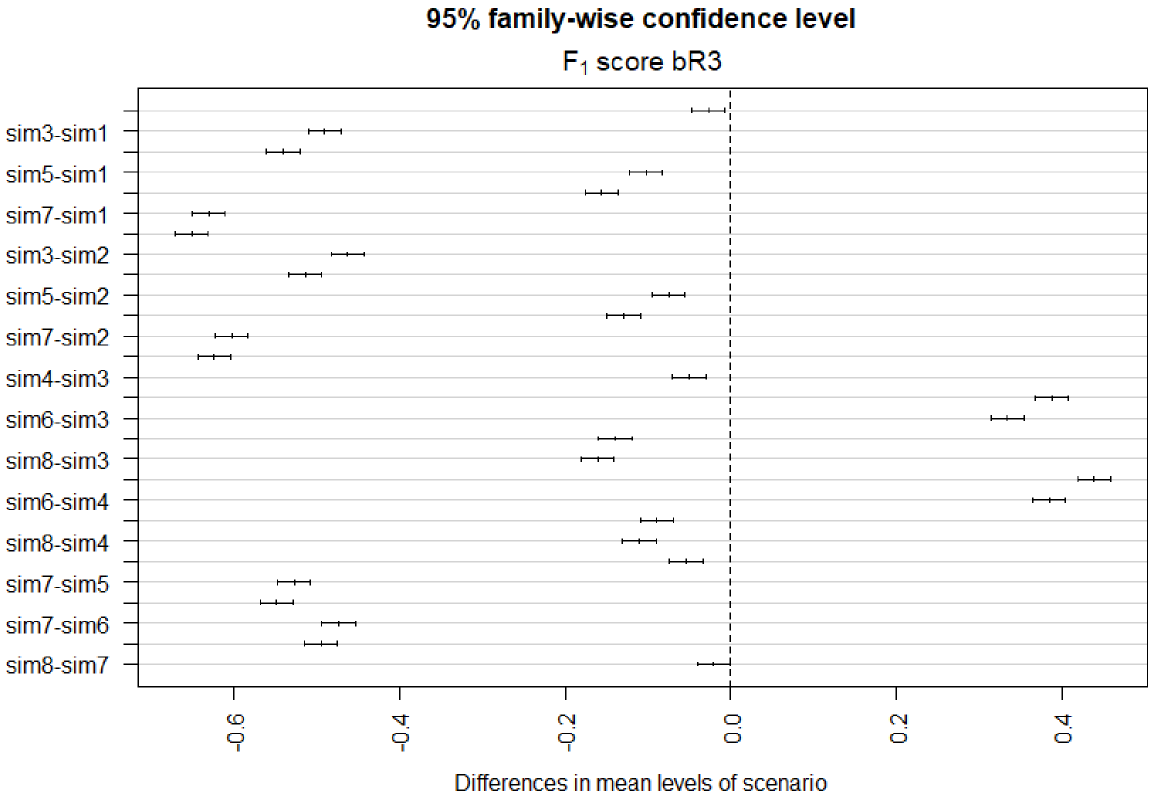


**S4 Fig. Pairwise comparison between $F_1$ scores, averaged across ten replicates, for the BayesR region-wide model (bR3) in different simulation scenarios for the quantitative phenotypes.** Simulation scenarios "sim1" to "sim8" are defined by combinations of simulated parameters in the following format ($h^2$_ $\pi$ _GA). sim1: 0.3_0.001_GA1, sim2: 0.3_0.001_GA2, sim3: 0.3_0.01_GA1, sim4: 0.3_0.01_GA2, sim5: 0.1_0.001_GA1, sim6: 0.1_0.001_GA2, sim7: 0.1_0.01_GA1, and sim8: 0.1_0.01_GA2. The dotted vertical line at the zero point on the x-axis indicates whether the average $F_1$ scores are significantly different.

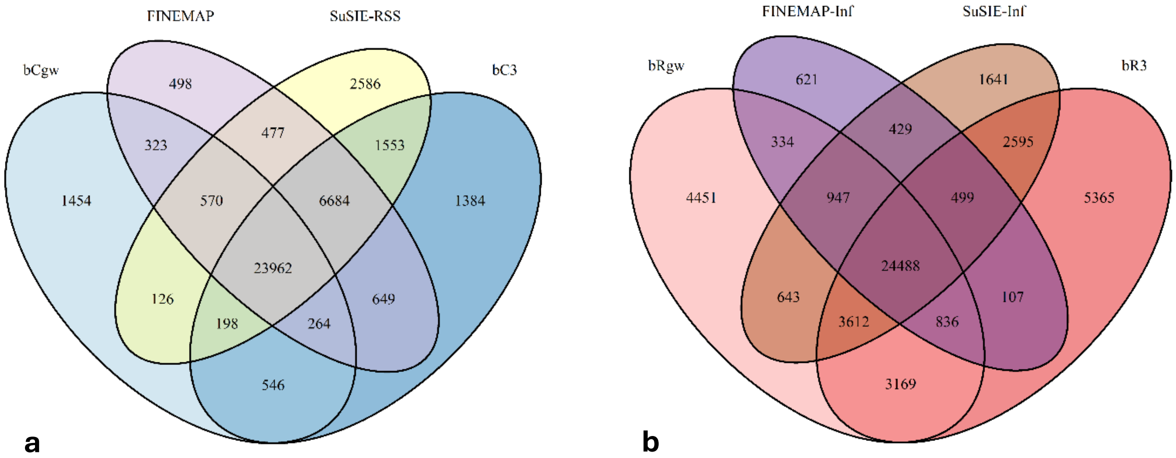


**S5 Fig. Venn-diagram for the common true positive credible sets summed across the eight simulation scenarios for the quantitative phenotypes**. **a.** Among BayesC genome-wide model (bCgw), BayesC region-wide model (bCgw), FINEMAP and SuSIE-RSS. **b.** Among BayesR genome-wide model (bRgw), BayesR region-wide model (bRgw), FINEMAP-Inf, and SuSIE-Inf.

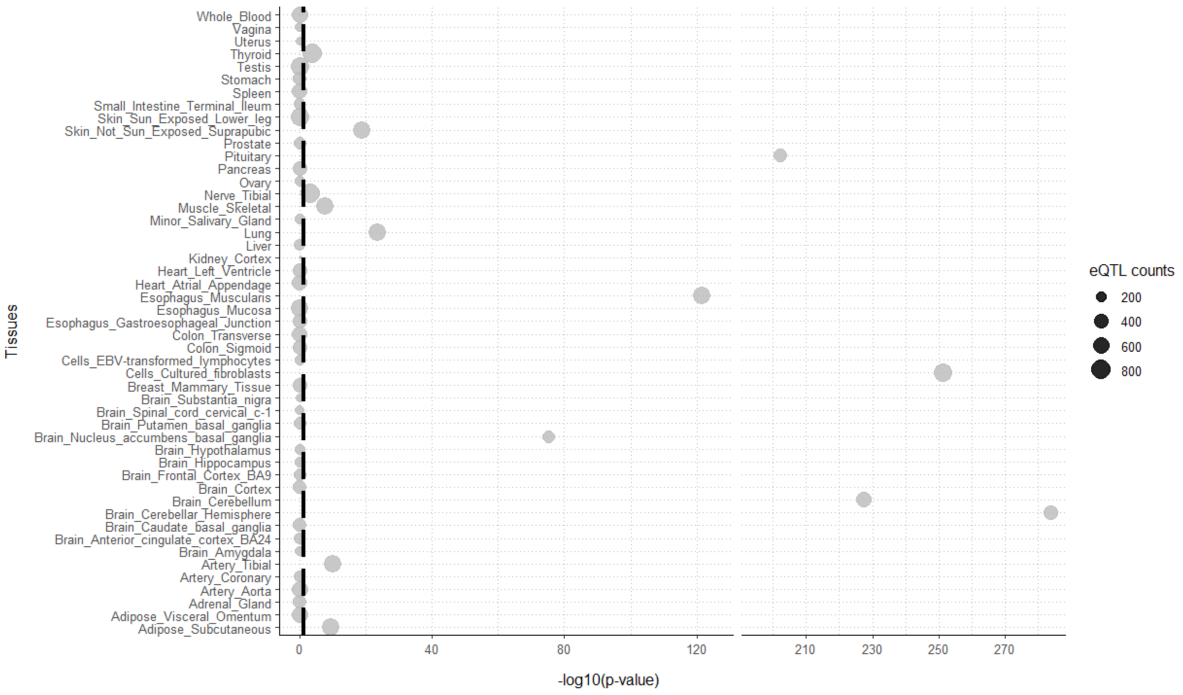


**S6 Fig. Significance values for enrichment analysis of tissue-specific eQTLs.** The black dashed line represents the significance cut-off (p-value < 0.05). The size of the points corresponds to the number of eQTLs (eQTL counts) in the tissue.
