## Supplementary Tables S1-S3 for "Evaluation of Bayesian Linear Regression Models as a Fine Mapping tool"

**S1 Table. Values for the parameters such as heritability, proportion of causal genetic variants, genetic architecture, and prevalence leading to different simulation scenarios for the quantitative and the binary phenotypes.**

| Quantitative simulation scenarios | Heritability ($h^2$) | Proportion of causal genetic variants (pi) | Genetic Architecture (GA) | Prevalence (PV) | Binary simulation scenarios |
|---|---|---|---|---|---|
| sim1 | 0.3 | 0.001 | GA1 | 0.05 | sim1 |
| | | | | 0.15 | sim2 |
| sim2 | 0.3 | 0.001 | GA2 | 0.05 | sim3 |
| | | | | 0.15 | sim4 |
| sim3 | 0.3 | 0.01 | GA1 | 0.05 | sim5 |
| | | | | 0.15 | sim6 |
| sim4 | 0.3 | 0.01 | GA2 | 0.05 | sim7 |
| | | | | 0.15 | sim8 |
| sim5 | 0.1 | 0.001 | GA1 | 0.05 | sim9 |
| | | | | 0.15 | sim10 |
| sim6 | 0.1 | 0.001 | GA2 | 0.05 | sim11 |
| | | | | 0.15 | sim12 |
| sim7 | 0.1 | 0.01 | GA1 | 0.05 | sim13 |
| | | | | 0.15 | sim14 |
| sim8 | 0.1 | 0.01 | GA2 | 0.05 | sim15 |
| | | | | 0.15 | sim16 |

**S2 Table. List of genes from Type two diabetes' credible sets, along with their position (chromosome, start and stop position), posterior inclusion probability (PIP), genetic variance (GenVar), and Overlap.** The Overlap column indicates if a gene is also located by the study Mahajan et al. 2018.

| Ensembl Gene ID | Symbol | Chr | Start | Stop | PIP | GenVar | Overlap |
|---|---|---|---|---|---|---|---|
| ENSG00000148737 | TCF7L2 | 10 | 114675346 | 114937385 | 2.8132 | 0.035914 | YES |
| ENSG00000118971 | CCND2 | 12 | 4347985 | 4424229 | 3.7104 | 0.011815 | YES |
| ENSG00000053918 | KCNQ1 | 11 | 2431014 | 2880204 | 2.8888 | 0.006736 | YES |
| ENSG00000073792 | IGF2BP2 | 3 | 185326662 | 185552635 | 0.9996 | 0.005882 | YES |
| ENSG00000140718 | FTO | 16 | 53703223 | 54165852 | 0.9996 | 0.004931 | YES |
| ENSG00000237941 | KCNQ1DN | 11 | 2856428 | 2903060 | 0.9996 | 0.004267 | NA |
| ENSG00000173175 | ADCY5 | 3 | 122966184 | 123178318 | 0.9996 | 0.003231 | YES |
| ENSG00000153814 | JAZF1 | 7 | 27835354 | 28230315 | 0.9996 | 0.003214 | YES |
| ENSG00000234336 | JAZF1-AS1 | 7 | 28185091 | 28293390 | 0.9996 | 0.003214 | NA |
| ENSG00000109501 | WFS1 | 4 | 6236816 | 6314892 | 0.9756 | 0.003122 | YES |
| ENSG00000074211 | PPP2R2C | 4 | 6287329 | 6575282 | 0.9756 | 0.003122 | NA |
| ENSG00000145996 | CDKAL1 | 6 | 20500048 | 21242620 | 0.838 | 0.003111 | YES |
| ENSG00000132170 | PPARG | 3 | 12293966 | 12485844 | 0.9996 | 0.003024 | YES |
| ENSG00000125746 | EML2 | 19 | 46075564 | 46158860 | 0.9952 | 0.002153 | NA |
| ENSG00000010310 | GIPR | 19 | 46136617 | 46196379 | 0.9952 | 0.002153 | YES |
| ENSG00000125743 | SNRPD2 | 19 | 46155919 | 46205777 | 0.9952 | 0.002153 | NA |
| ENSG00000180176 | TH | 11 | 2150304 | 2203045 | 0.9996 | 0.002076 | NA |
| ENSG00000130204 | TOMM40 | 19 | 45358937 | 45416831 | 0.9996 | 0.00198 | YES |
| ENSG00000130203 | APOE | 19 | 45374101 | 45422561 | 0.9996 | 0.00198 | YES |
| ENSG00000130208 | APOC1 | 19 | 45382675 | 45432557 | 0.9996 | 0.00198 | NA |
| ENSG00000214855 | APOC1P1 | 19 | 45395193 | 45444566 | 0.9996 | 0.00198 | NA |
| ENSG00000267467 | APOC4 | 19 | 45410911 | 45461996 | 0.9996 | 0.00198 | NA |
| ENSG00000084734 | GCKR | 2 | 27684734 | 27755993 | 0.9988 | 0.001845 | YES |
| ENSG00000108175 | ZMIZ1 | 10 | 80793869 | 81085599 | 0.8776 | 0.00183 | YES |
| ENSG00000186635 | ARAP1 | 11 | 72361117 | 72514533 | 0.9944 | 0.001811 | YES |
| ENSG00000214530 | STARD10 | 11 | 72430935 | 72514533 | 0.9944 | 0.001811 | NA |
| ENSG00000159217 | IGF2BP1 | 17 | 47040024 | 47142930 | 0.996 | 0.001647 | NA |
| ENSG00000064655 | EYA2 | 20 | 45488529 | 45827115 | 0.9828 | 0.001553 | YES |
| ENSG00000134250 | NOTCH2 | 1 | 120420048 | 120622021 | 0.9996 | 0.001519 | YES |
| ENSG00000182247 | UBE2E2 | 3 | 23210370 | 23643220 | 0.962 | 0.001439 | YES |
| ENSG00000108753 | HNF1B | 17 | 36011707 | 36115166 | 1.0044 | 0.001353 | YES |
| ENSG00000152527 | PLEKHH2 | 2 | 43829540 | 44004860 | 0.7956 | 0.001328 | NA |
| ENSG00000154122 | ANKH | 5 | 14670509 | 14881275 | 0.9428 | 0.001307 | YES |
| ENSG00000171791 | BCL2 | 18 | 60755789 | 60996360 | 0.9976 | 0.001278 | NA |
| ENSG00000204469 | PRRC2A | 6 | 31553935 | 31615514 | 0.8444 | 0.001243 | NA |
| ENSG00000204463 | BAG6 | 6 | 31571894 | 31630241 | 0.8444 | 0.001243 | NA |
| ENSG00000204444 | APOM | 6 | 31585219 | 31635943 | 0.8444 | 0.001243 | NA |

| | | | | | | | |
|---|---|---|---|---|---|---|---|
| ENSG00000204439 | C6orf47 | 6 | 31591198 | 31638178 | 0.8444 | 0.001243 | NA |
| ENSG00000204438 | GPANK1 | 6 | 31594181 | 31643522 | 0.8444 | 0.001243 | NA |
| ENSG00000204435 | CSNK2B | 6 | 31598037 | 31647666 | 0.8444 | 0.001243 | NA |
| ENSG00000240053 | LY6G5B | 6 | 31602967 | 31651247 | 0.8444 | 0.001243 | NA |
| ENSG00000204428 | LY6G5C | 6 | 31609813 | 31661708 | 0.8444 | 0.001243 | NA |
| ENSG00000130749 | ZC3H4 | 19 | 47532517 | 47626543 | 0.9092 | 0.001229 | YES |
| ENSG00000082438 | COBLL1 | 2 | 165476253 | 165710111 | 1.002 | 0.001205 | YES |
| ENSG00000160360 | GPSM1 | 9 | 139217089 | 139263891 | 0.8908 | 0.001193 | YES |
| ENSG00000213221 | DNLZ | 9 | 139219025 | 139267838 | 0.8908 | 0.001193 | NA |
| ENSG00000187796 | CARD9 | 9 | 139221567 | 139277875 | 0.8908 | 0.001193 | NA |
| ENSG00000165684 | SNAPC4 | 9 | 139235370 | 139303049 | 0.8908 | 0.001193 | NA |
| ENSG00000152601 | MBNL1 | 3 | 151926894 | 152193063 | 0.9436 | 0.001128 | YES |
| ENSG00000164756 | SLC30A8 | 8 | 117927563 | 118198669 | 0.6192 | 0.001113 | YES |
| ENSG00000185122 | HSF1 | 8 | 145491340 | 145548143 | 0.8608 | 0.001095 | NA |
| ENSG00000185000 | DGAT1 | 8 | 145505126 | 145559062 | 0.8608 | 0.001095 | NA |
| ENSG00000170616 | SCRT1 | 8 | 145519687 | 145569849 | 0.8608 | 0.001095 | NA |
| ENSG00000214597 | TMEM249 | 8 | 145541424 | 145587615 | 0.8608 | 0.001095 | NA |
| ENSG00000185803 | SLC52A2 | 8 | 145543014 | 145594363 | 0.8608 | 0.001095 | NA |
| ENSG00000182325 | FBXL6 | 8 | 145544720 | 145592444 | 0.8608 | 0.001095 | NA |
| ENSG00000153815 | CMIP | 16 | 81443859 | 81754625 | 0.8908 | 0.001089 | YES |
| ENSG00000165917 | RAPSN | 11 | 47424400 | 47480403 | 0.9216 | 0.001086 | NA |
| ENSG00000149187 | CELF1 | 11 | 47452744 | 47596789 | 0.9216 | 0.001086 | YES |
| ENSG00000008196 | TFAP2B | 6 | 50751885 | 50824804 | 0.8944 | 0.001066 | YES |
| ENSG00000060303 | RPS17P5 | 6 | 50790633 | 50835180 | 0.8944 | 0.001066 | NA |
| ENSG00000124783 | SSR1 | 6 | 7234172 | 7357333 | 0.9012 | 0.000989 | NA |
| ENSG00000185104 | FAF1 | 1 | 50870261 | 51433687 | 0.8144 | 0.000979 | YES |
| ENSG00000124782 | RREB1 | 6 | 7073433 | 7261985 | 0.7836 | 0.000972 | YES |
| ENSG00000157895 | C12orf43 | 12 | 121405288 | 121463392 | 1.7392 | 0.000955 | NA |
| ENSG00000135100 | HNF1A | 12 | 121381353 | 121450165 | 1.528 | 0.000888 | YES |
| ENSG00000151465 | CDC123 | 10 | 12203296 | 12302357 | 0.6208 | 0.000847 | YES |
| ENSG00000152270 | PDE3B | 11 | 14630370 | 14901495 | 0.8556 | 0.000816 | YES |
| ENSG00000186104 | CYP2R1 | 11 | 14865551 | 14923717 | 0.8556 | 0.000816 | NA |
| ENSG00000102908 | NFAT5 | 16 | 69564247 | 69747925 | 0.6068 | 0.00078 | YES |
| ENSG00000078237 | TIGAR | 12 | 4395865 | 4472276 | 0.9008 | 0.000765 | NA |
| ENSG00000134640 | MTNR1B | 11 | 92667946 | 92728050 | 0.88 | 0.000764 | YES |
| ENSG00000087448 | KLHL42 | 12 | 27898264 | 27965869 | 0.858 | 0.000737 | NA |
| ENSG00000061794 | MRPS35 | 12 | 27828938 | 27919145 | 0.7664 | 0.000729 | NA |
| ENSG00000205693 | MANSC4 | 12 | 27881205 | 27932562 | 0.7664 | 0.000729 | NA |
| ENSG00000029534 | ANK1 | 8 | 41476003 | 41764243 | 1.0356 | 0.000723 | YES |
| ENSG00000130584 | ZBTB46 | 20 | 62340115 | 62472332 | 0.8972 | 0.000718 | YES |
| ENSG00000183734 | ASCL2 | 11 | 2255018 | 2301727 | 0.8788 | 0.00061 | NA |
| ENSG00000110665 | C11orf21 | 11 | 2282077 | 2334039 | 0.8788 | 0.00061 | NA |
| ENSG00000004534 | RBM6 | 3 | 49943200 | 50147188 | 0.676 | 0.00056 | YES |
| ENSG00000145730 | PAM | 5 | 102054834 | 102376521 | 0.9092 | 0.000543 | YES |
| ENSG00000196781 | TLE1 | 9 | 84163625 | 84313802 | 0.9524 | 0.000543 | YES |
| ENSG00000168928 | CTRB2 | 16 | 75202998 | 75250862 | 0.9724 | 0.000465 | NA |

| | | | | | | | |
|---|---|---|---|---|---|---|---|
| ENSG00000168925 | CTRB1 | 16 | 75217943 | 75268523 | 0.9724 | 0.000465 | NA |
| ENSG00000050820 | BCAR1 | 16 | 75227969 | 75311870 | 0.9724 | 0.000465 | YES |
| ENSG00000171017 | LRRC8E | 19 | 7918993 | 7976698 | 0.9032 | 0.000464 | NA |
| ENSG00000076984 | MAP2K7 | 19 | 7933748 | 7988117 | 0.9032 | 0.000464 | YES |
| ENSG00000104976 | SNAPC2 | 19 | 7950288 | 7997870 | 0.9032 | 0.000464 | NA |
| ENSG00000178531 | CTXN1 | 19 | 7954573 | 8000727 | 0.9032 | 0.000464 | NA |
| ENSG00000104980 | TIMM44 | 19 | 7957481 | 8018029 | 0.9032 | 0.000464 | NA |
| ENSG00000165066 | NKX6-3 | 8 | 41467827 | 41517860 | 0.5012 | 0.000462 | NA |
| ENSG00000172575 | RASGRP1 | 15 | 38745465 | 38867413 | 0.8356 | 0.000444 | YES |
| ENSG00000119772 | DNMT3A | 2 | 25420865 | 25574987 | 0.7504 | 0.00039 | NA |
| ENSG00000158186 | MRAS | 3 | 138033181 | 138134327 | 0.9448 | 0.000343 | NA |
| ENSG00000135114 | OASL | 12 | 121423167 | 121487014 | 0.5676 | 0.00024 | NA |
| ENSG00000002822 | MAD1L1 | 7 | 1820538 | 2282612 | 0.8344 | 0.000202 | NA |
| ENSG00000149948 | HMGA2 | 12 | 66182924 | 66369804 | 2.0908 | 0.000199 | YES |
| ENSG00000009335 | UBE3C | 7 | 156896804 | 157071972 | 0.6556 | 0.000182 | NA |
| ENSG00000181322 | NME9 | 3 | 137945606 | 138057915 | 0.398 | 0.000179 | NA |
| ENSG00000072110 | ACTN1 | 14 | 69306149 | 69456011 | 0.364 | 0.000147 | NA |
| ENSG00000117707 | PROX1 | 1 | 214121995 | 214224428 | 1.2612 | 0.000144 | YES |
| ENSG00000158220 | ESYT3 | 3 | 138118488 | 138209748 | 0.304 | 0.0001 | NA |
| ENSG00000198373 | WWP2 | 16 | 69761563 | 69984145 | 0.26 | 8.84E-05 | NA |
| ENSG00000157322 | CLEC18A | 16 | 69950530 | 70007531 | 0.26 | 8.84E-05 | NA |
| ENSG00000003756 | RBM5 | 3 | 50092226 | 50165563 | 0.2088 | 8.69E-05 | NA |
| ENSG00000011523 | CEP68 | 2 | 65249014 | 65324089 | 0.8148 | 6.31E-05 | YES |
| ENSG00000138069 | RAB1A | 2 | 65262906 | 65367056 | 0.8148 | 6.31E-05 | NA |
| ENSG00000204929 | LOC101927533 | 2 | 65629079 | 66321554 | 1.6056 | 5.84E-05 | NA |
| ENSG00000138101 | DTNB | 2 | 25565907 | 25906274 | 0.2116 | 5.76E-05 | YES |
| ENSG00000198369 | SPRED2 | 2 | 65503110 | 65668516 | 0.7388 | 5.36E-05 | NA |
| ENSG00000141699 | RETREG3 | 17 | 40696679 | 40772503 | 0.506 | 3.56E-05 | NA |
| ENSG00000115970 | THADA | 2 | 43358820 | 43833039 | 0.1388 | 3.39E-05 | YES |
| ENSG00000176095 | IP6K1 | 3 | 49727101 | 49833904 | 0.1452 | 2.98E-05 | NA |
| ENSG00000147889 | CDKN2A | 9 | 21933125 | 22004322 | 0.1764 | 2.87E-05 | YES |
| ENSG00000147883 | CDKN2B | 9 | 21968155 | 22019156 | 0.1764 | 2.87E-05 | NA |
| ENSG00000131462 | TUBG1 | 17 | 40726734 | 40777251 | 0.3312 | 2.22E-05 | NA |
| ENSG00000183260 | ABHD16B | 20 | 62458395 | 62504309 | 0.1368 | 1.86E-05 | NA |
| ENSG00000101150 | TPD52L2 | 20 | 62461655 | 62532894 | 0.1368 | 1.86E-05 | NA |
| ENSG00000181751 | MACIR | 5 | 102559407 | 102624161 | 0.1456 | 1.78E-05 | NA |
| ENSG00000267132 | HMGB3P27 | 17 | 40766546 | 40810228 | 0.2328 | 1.58E-05 | NA |
| ENSG00000131470 | PSMC3IP | 17 | 40689455 | 40738610 | 0.2184 | 1.52E-05 | NA |
| ENSG00000176349 | RP11-69E11.4 | 7 | 1843294 | 1899479 | 0.11 | 1.41E-05 | NA |
| ENSG00000068120 | COASY | 17 | 40678723 | 40728229 | 0.1748 | 1.34E-05 | NA |
| ENSG00000108788 | MLX | 17 | 40684176 | 40735077 | 0.1748 | 1.34E-05 | YES |
| ENSG00000108784 | NAGLU | 17 | 40653534 | 40706273 | 0.1252 | 1.09E-05 | NA |
| ENSG00000108786 | HSD17B1 | 17 | 40667572 | 40716520 | 0.1252 | 1.09E-05 | NA |
| ENSG00000037042 | TUBG2 | 17 | 40776904 | 40828929 | 0.1216 | 9.32E-06 | NA |
| ENSG00000068137 | PLEKHH3 | 17 | 40785272 | 40838829 | 0.0828 | 7.77E-06 | NA |
| ENSG00000127603 | MACF1 | 1 | 39512153 | 39962593 | 3.198 | 7.63E-06 | YES |

| | | | | | | | |
|---|---|---|---|---|---|---|---|
| ENSG00000033627 | ATP6V0A1 | 17 | 40576162 | 40684176 | 0.0764 | 2.52E-06 | NA |
| ENSG00000177469 | CAVIN1 | 17 | 40519890 | 40584708 | 0.0744 | 2.18E-06 | NA |
| ENSG00000174099 | MSRB3 | 12 | 65638037 | 65891257 | 0.7632 | 2.17E-06 | NA |
| ENSG00000175832 | ETV4 | 17 | 41570427 | 41666423 | 0.5536 | 1.07E-06 | NA |
| ENSG00000163909 | HEYL | 1 | 40055180 | 40115514 | 0.3552 | 9.60E-07 | NA |
| ENSG00000188554 | NBR1 | 17 | 41287708 | 41372775 | 0.1744 | 4.79E-07 | NA |
| ENSG00000214114 | MYCBP | 1 | 39293721 | 39357157 | 0.5524 | 4.20E-07 | NA |
| ENSG00000131233 | GJA9 | 1 | 39295433 | 39357157 | 0.5524 | 4.20E-07 | NA |
| ENSG00000158315 | RHBDL2 | 1 | 39316517 | 39417469 | 0.6292 | 4.15E-07 | NA |
| ENSG00000184988 | TMEM106A | 17 | 41329010 | 41381753 | 0.0908 | 4.11E-07 | NA |
| ENSG00000090621 | PABPC4 | 1 | 39991588 | 40052108 | 0.3188 | 3.93E-07 | NA |
| ENSG00000188825 | LINC00910 | 17 | 41412250 | 41476397 | 0.2392 | 3.23E-07 | NA |
| ENSG00000175906 | ARL4D | 17 | 41442419 | 41488496 | 0.2116 | 3.11E-07 | NA |
| ENSG00000067596 | DHX8 | 17 | 41526526 | 41631615 | 0.3888 | 2.76E-07 | NA |
| ENSG00000116954 | RRAGC | 1 | 39269459 | 39335493 | 0.2944 | 2.59E-07 | NA |
| ENSG00000182109 | NA | 1 | 39952956 | 40021644 | 0.3248 | 1.46E-07 | NA |
| ENSG00000182197 | EXT1 | 8 | 118771912 | 119133925 | 0.0236 | 1.38E-07 | NA |
| ENSG00000012048 | BRCA1 | 17 | 41161500 | 41287151 | 0.2728 | 1.26E-07 | NA |
| ENSG00000117000 | RLF | 1 | 40592848 | 40716509 | 0.3772 | 1.16E-07 | NA |
| ENSG00000198496 | NBR2 | 17 | 41243190 | 41315364 | 0.1496 | 8.69E-08 | NA |
| ENSG00000117010 | ZNF684 | 1 | 40962617 | 41023763 | 0.5148 | 6.88E-08 | NA |
| ENSG00000139746 | RBM26 | 13 | 79851031 | 79990417 | 0.446 | 5.53E-08 | NA |
| ENSG00000102471 | NDFIP2 | 13 | 80020616 | 80139875 | 0.1636 | 4.98E-08 | NA |
| ENSG00000131236 | CAP1 | 1 | 40470939 | 40548306 | 0.8172 | 4.46E-08 | NA |
| ENSG00000188800 | TMCO2 | 1 | 40677050 | 40727279 | 0.1344 | 4.31E-08 | NA |
| ENSG00000183682 | BMP8A | 1 | 39922426 | 40001489 | 0.358 | 4.08E-08 | NA |
| ENSG00000084073 | ZMPSTE24 | 1 | 40690467 | 40769783 | 0.1836 | 3.57E-08 | NA |
| ENSG00000164002 | EXO5 | 1 | 40939613 | 40992208 | 0.274 | 2.86E-08 | NA |
| ENSG00000131238 | PPT1 | 1 | 40503635 | 40573134 | 0.5788 | 2.51E-08 | NA |
| ENSG00000068079 | IFI35 | 17 | 41124108 | 41174175 | 0.0844 | 2.27E-08 | NA |
| ENSG00000108828 | VAT1 | 17 | 41131645 | 41187009 | 0.0864 | 2.06E-08 | NA |
| ENSG00000198863 | RUNDC1 | 17 | 41098108 | 41155531 | 0.0684 | 1.96E-08 | NA |
| ENSG00000108825 | PTGES3L-AARSD1 | 17 | 41067670 | 41141971 | 0.0528 | 1.79E-08 | NA |
| ENSG00000267060 | PTGES3L | 17 | 41085683 | 41141971 | 0.0528 | 1.79E-08 | NA |
| ENSG00000108830 | RND2 | 17 | 41143775 | 41193910 | 0.0872 | 1.63E-08 | NA |
| ENSG00000187815 | ZFP69 | 1 | 40908385 | 40971820 | 0.1584 | 1.62E-08 | NA |
| ENSG00000049089 | COL9A2 | 1 | 40732348 | 40793456 | 0.0916 | 1.49E-08 | NA |
| ENSG00000131469 | RPL27 | 17 | 41116836 | 41163974 | 0.06 | 1.45E-08 | NA |
| ENSG00000237888 | AC087650.1 | 17 | 41518927 | 41563997 | 0.0984 | 1.25E-08 | NA |
| ENSG00000266967 | AARSD1 | 17 | 41067670 | 41124108 | 0.0316 | 9.52E-09 | NA |
| ENSG00000267151 | MIR2117HG | 17 | 41487103 | 41538446 | 0.0736 | 8.30E-09 | NA |
| ENSG00000145725 | PPIP5K2 | 5 | 102420854 | 102558205 | 0.1308 | 5.44E-09 | NA |
| ENSG00000187801 | ZFP69B | 1 | 40880804 | 40939387 | 0.0244 | 3.93E-09 | NA |
| ENSG00000205359 | SLCO6A1 | 5 | 101673587 | 101843903 | 0.0692 | 2.88E-09 | YES |
| ENSG00000145723 | GIN1 | 5 | 102386985 | 102465231 | 0.0468 | 2.11E-09 | NA |

| | | | | | | | |
|---|---|---|---|---|---|---|---|
| ENSG00000116985 | BMP8B | 1 | 40188113 | 40264168 | 0.3564 | 1.81E-09 | NA |
| ENSG00000043514 | TRIT1 | 1 | 40271768 | 40358726 | 0.3544 | 1.51E-09 | NA |
| ENSG00000116990 | MYCL | 1 | 40326106 | 40377735 | 0.1788 | 1.01E-09 | NA |
| ENSG00000168389 | MFSD2A | 1 | 40385934 | 40445199 | 0.0116 | 6.66E-10 | NA |
| ENSG00000173930 | SLCO4C1 | 5 | 101534745 | 101642159 | 0.038 | 5.00E-10 | NA |
| ENSG00000198754 | OXCT2 | 1 | 40200894 | 40246925 | 0.0476 | 2.57E-10 | NA |
| ENSG00000084072 | PPIE | 1 | 40122939 | 40238449 | 0.0272 | 2.40E-10 | NA |

***YES**: overlapped with Mahajan et al. 2018; **NA**: not overlapped with Mahajan et al. 2018; **PIP**: sum of posterior inclusion probability of SNPs in a gene; **GenVar**: Genetic variance explained by a gene.*

**S3 Table. Top 30 diseases, from DISEASE database, associated with Type two diabetes' gene sets.**

| Number | Disease | P-value | Remarks |
|---|---|---|---|
| 1 | Maturity-onset diabetes of the young type 4 | 4.44E-16 | |
| 2 | Prediabetes syndrome | 6.66E-16 | |
| 3 | Gestational diabetes | 2.89E-15 | |
| 4 | ICD10:O24 | 2.89E-15 | Diabetes mellitus in pregnancy |
| 5 | ICD10:O2 | 3.44E-15 | Other abnormal products of conception |
| 6 | Permanent neonatal diabetes mellitus | 3.10E-14 | |
| 7 | Pancreatic agenesis | 1.28E-13 | |
| 8 | ICD10:E11 | 1.71E-13 | Type two diabetes (T2D) |
| 9 | Type 2 diabetes mellitus | 1.71E-13 | |
| 10 | Wolfram syndrome | 2.02E-13 | |
| 11 | Maturity-onset diabetes of the young type 9 | 6.05E-13 | |
| 12 | Maturity-onset diabetes of the young type 2 | 7.09E-13 | |
| 13 | Maturity-onset diabetes of the young type 10 | 1.66E-12 | |
| 14 | ICD10:E1 | 3.36E-12 | Iodine-deficiency-related thyroid disorders and allied conditions |
| 15 | Wolfram syndrome 1 | 3.41E-12 | |
| 16 | Diabetes mellitus | 3.78E-12 | |
| 17 | ICD10:E14 | 3.78E-12 | Unspecified T2D |
| 18 | Maturity-onset diabetes of the young type 1 | 4.60E-12 | |
| 19 | Hyperinsulinism | 5.51E-12 | |
| 20 | ICD10:E161 | 5.51E-12 | MS: E16.1 - Other hypoglycemia |
| 21 | Insulinoma | 5.56E-12 | |
| 22 | ICD10:M06 | 6.39E-12 | Other Rheumatoid Arthritis (RA) |
| 23 | ICD10:M069 | 6.39E-12 | RA |
| 24 | Rheumatoid arthritis | 6.39E-12 | |
| 25 | ICD10:M05 | 6.49E-12 | Felty syndrome (RA with splenoadenomegaly and leukopenia) |
| 26 | ICD10:M0 | 9.86E-12 | Pyogenic arthritis |
| 27 | Hyperglycemia | 1.10E-11 | |
| 28 | Pancreatic cystadenoma | 1.27E-11 | |
| 29 | Glucose intolerance | 1.64E-11 | |
| 30 | Transient neonatal diabetes mellitus | 1.65E-11 | |
