## Supplementary material for "Evaluation of Bayesian Linear Regression Models as a Fine Mapping tool": S1 Text

### S1. Design of multiple credible sets in a fine-mapping region for the UKB phenotypes

To explore the presence of multiple CSs within a fine-mapped region (Figure S1), we followed these steps until all SNPs were evaluated:

1. We assessed the region for SNPs with a $PIP_{SNP}$ of at least 0.80.
    a. If a SNP (SNPs) met the criterion $PIP_{SNP}$ >= 0.80 a CS (CSs) is formed. SNPs used in this step were then excluded from further analysis before moving onto the second step. The analysis continued to the second step if no SNPs met the initial PIP criterion.
2. In the second step, we identified the SNP (for example SNP-A) with the highest PIP. We then searched for other SNPs in LD with SNP-A that met the LD criterion.
    a. If no SNPs were in LD with SNP-A (meeting the LD criterion), SNP-A was removed from the region without forming a CS. If there were SNPs in LD with SNP-A (SNP-A-set1), we evaluated if the cumulative PIP of this set (SNP-A-set1) in descending order or its' subset met the criterion $PIP_{cums-set}$ >= 0.80).
    b. If the set failed to meet the PIP criterion, SNP-A was excluded. If the set met the criterion, it was recognized as a CS, and the SNPs were removed from the region. This could result in a CS such as SNP-A-set1 or a subset like SNP-A-set2.
3. The analysis continued from the second step until no SNPs remained for analysis.
